# Informational and methodological differences in regional structure-function coupling in modeling approaches

**DOI:** 10.1101/2025.10.26.684597

**Authors:** Yihan Zhang

**Affiliations:** School of Science, China Pharmaceutical University, Nanjing 211198, China

**Keywords:** structural connectivity, functional connectivity, structure–function coupling, indirect connections, approach comparison, graph neural networks

## Abstract

Structural connectivity in the human brain constrains functional connectivity, which in turn reshapes structural connectivity, resulting in complex coupling relationships. Numerous approaches exist for quantifying structure-function coupling, and different approaches exhibit systematic differences in their estimations, but the sources and specific manifestations of these differences remain unclear. In this study, starting from the perspective of indirect connection information and using the correlational approach as a baseline, we systematically compared the differences among four modeling approaches (multivariate linear regression, multilayer perceptron, predictive graph neural network and self-supervised graph neural network) in the calculation of regional structure-function coupling by defining the informational and methodological differences. Our results show that indirect connections have a small effect on the regression approach and the multilayer perceptron, while a larger effect on the graph neural networks. Furthermore, we observed that, in predicting functional connectivity, the prediction of direct connections of brain regions with higher T1w/T2w myelination indices is more significantly influenced by indirect connections. At the level of structure-function coupling, the right-hemispheric dorsal attention network is least affected by indirect connections, while the orbito-affective network is most significantly affected. These findings reveal the mechanisms of information usage behind different approaches in the calculation of structure-function coupling, providing a basis for future research to select appropriate approaches based on the structural and functional network characteristics of the brain.

## 1 Introduction

Understanding the correspondence between the human brain’s anatomical structure and physiological function is one of the main goals of neuroscience. The structural characteristics of the brain are often described in terms of white matter connections between brain regions, which form structural connectomes with complex topological properties that strongly constrain neuronal activity (Park & Friston, 2013). Meanwhile, neuronal activity can remodel structural connections through neuromodulation and neuroplasticity processes (Bargmann, 2012; Froemke, 2015) to adapt to different cognitive tasks and complex external environments. A commonly used method for inferring structural connectivity is tractography based on diffusion-weighted imaging (DWI) (Catani et al., 2002; Sotiropoulos & Zalesky, 2019), which can quantify the number of white matter fiber tracts between brain regions, thus measuring the strength of structural connectivity. In contrast, functional connectivity can be measured by calculating the statistical correlation of neural activity time series from functional magnetic resonance imaging (fMRI).

Functional connectivity is constrained by structural connectivity on a large scale and is positively correlated with it. However, owing to the complexity of multisynaptic interactions and the presence of indirect connections, this correspondence is not perfect (Damoiseaux & Greicius, 2009; Honey et al., 2009; Suárez et al., 2020). To further quantify the structure-function coupling (SFC) of different brain regions, the most commonly used approach is the correlational approach, which defines the SFC of each brain region as the correlation coefficient between its structural and functional connectivity vectors with other brain regions (Cocchi et al., 2014). The second is the modeling approach, which mostly uses the paradigm of predicting functional connectivity from structural connectivity (or other prediction tasks). This includes both computational and statistical models for direct modeling (Finger et al., 2016; Vázquez-Rodríguez et al., 2019) and data-driven deep learning models (Chen et al., 2024; Li et al., 2022; Sarwar et al., 2021). These approaches effectively address the extent to which functional connectivity depends on structural connectivity and demonstrate the imperfect coupling between the two.

In general, SFC is stronger in the primary sensorimotor cortex and weaker in the association cortex. This conclusion has been validated in multiple studies using various approaches and perspectives (Baum et al., 2022; Collins et al., 2024; M. F. Glasser & Essen, 2011; Valk et al., 2022). However, the specific distribution of SFC is influenced by a variety of factors such as age (Esfahlani et al., 2022; Liu et al., 2024), gender (Zhao et al., 2021), individual differences (Chen et al., 2024; Gu et al., 2021), and cognitive state (Hermundstad et al., 2014). Even with the same sample, different approaches often yield different estimates. For example, the SFC calculated by deep learning models is higher than that calculated by the correlational approach and biophysical models (Chen et al., 2024; Sarwar et al., 2021). In short, no approach has yet emerged that can completely and accurately capture structure-function coupling. Each approach has different degrees of assumptions. These different assumptions allow for the capture of coupling at different levels, thus enabling the solving of different tasks such as connectivity prediction (Chen et al., 2024; Sarwar et al., 2021), disease diagnosis (Sun et al., 2024; Wu et al., 2025), and cognitive state classification (Griffa et al., 2022). We can simply consider these assumptions as the systematic differences between different approaches, determining how connectivity information is used and how the correspondence between structure and function is estimated. However, due to the numerous approaches for calculating SFC, it is difficult to uniformly answer how these systematic differences are manifested.

One approach is to compare approaches in specific scenarios, such as disease diagnosis (Lu et al., 2025; Wei et al., 2025), cognitive state classification (Kucukosmanoglu et al., 2024), and quantification of individual differences (Pan et al., 2025). This helps to intuitively compare different approaches and select the optimal approach for a specific task, but its limitation is that it only answers the differences between approaches within a specific scenario. Another approach is to compare models within a unified perspective, such as predictors (Esfahlani et al., 2022), eigendecompositions (Deslauriers-Gauthier et al., 2020), or a combination of multiple perspectives (Messé et al., 2015). This approach allows for a deeper comparison of the specific mechanisms by which different models learn the coupling between structure and function. However, its generalizability is also affected by the chosen perspective, thus most studies can only compare a specific type of model. Our study follows this second approach, attempting to explain the differences in the calculation of regional SFC by examining whether or not indirect connections are used.

The introduction of indirect connections makes regional SFC no longer local, but influenced by connectivity information from a broader region. Previous studies have shown that indirect connections help improve the model’s ability to predict functional connectivity (Esfahlani et al., 2022; Røge et al., 2017), and many models naturally consider indirect connections when predicting functional connectivity, especially the increasingly used deep learning models in recent years. However, the way different models utilize indirect connections and the extent to which they affect SFC estimation have not been systematically compared. For example, models that predict functional connectivity from structural connectivity implicitly assume that functional connectivity can be derived from structural connectivity (Chen et al., 2024; Sarwar et al., 2021), while other models directly calculate the coupling in the interaction between structure and function (Li et al., 2022; Xia et al., 2025), the types of indirect connections they rely on differ. Therefore, to investigate whether there are systematic differences in the sensitivity of different models to indirect connections, and whether such differences are brain region specific, we compared several common modeling approaches using the correlational approach as a baseline.

In this study, we systematically compared the differences in the calculation of regional SFC among four modeling approaches: regression approach (Vázquez-Rodríguez et al., 2019), multilayer perceptron (MLP) (Sarwar et al., 2021), predictive graph convolutional network (pGCN) (Chen et al., 2024) and self-supervised graph convolutional network (sGCN), and using the correlational approach (Baum et al., 2020; Gu et al., 2021; Honey et al., 2009; Liégeois et al., 2020) as a baseline. To this end, we defined the informational difference (caused by different combinations of connectivity information) and the methodological difference (difference that still exists under the same information combination), and separated the two by masking indirect connections. Our results show that indirect connections have a small effect on the regression approach and the MLP, but a larger effect on the pGCN and the sGCN. Furthermore, the informational difference resulting from indirect connections is specific to different cortical regions: at the predicted-connectivity level, its magnitude is positively correlated with T1w/T2w myelination index of the cortical region; at the SFC level, the informational difference is lowest in the right-hemispheric dorsal attention network and highest in the orbito-affective network. These findings reveal the mechanism of information usage behind different SFC calculation approaches, providing a basis for future research to select appropriate approaches based on specific brain regions or network characteristics. Figure 4 provides a schematic overview of this study.

## 2 Methods

### 2.1 Dataset

#### 2.1.1 Data sources

We obtained 3T resting-state function magnetic resonance (rs-fMRI) data from the WU-Minn Human Connectome Project (HCP) 1200 subjects data release (Essen et al., 2013), which had been preprocessed with ICA-FIX denoising. The rs-fMRI data comprise 1003 subjects (469 males, 22-36+ years old). All participants provided informed consent, approved by the Institutional Review Board at Washington University in St. Louis. All rs-fMRI data were acquired using a customized Siemens 3T Skyra scanner in four runs of approximately 15 minutes with 1200 frames each, coming from two sessions (each containing two runs) with opposite phase encoding directions: right-to-left (RL) in one run and left-to-right (LR) in the other. Other detailed descriptions of the HCP S1200 dataset and acquisition parameters can be found in the manual (WU-Minn HCP, 2017) and prior study (Essen et al., 2013).

Structural connectivity data were obtained from the publicly available resource *A Whole-Cortex Probabilistic Diffusion Tractography Connectome* (Rosen & Halgren, 2021). This dataset provides preprocessed structural connectivity matrices for 1065 HCP S1200 subjects (490 males, 22-36+ years old, 998 of which also had rs-fMRI data). The diffusion MRI data in the HCP S1200 dataset used by Rosen and Halgren (2021) were acquired in 6 runs of approximately 9 minutes and 50 seconds each. The 6 runs represented 3 different gradient tables with two opposite phase encoding directions (RL and LR) each. Each gradient table includes approximately 90 directions and 3 shells (b = 1000, 2000 and 3000 s/mm^2^). Other detailed acquisition parameters can be found in the manual (WU-Minn HCP, 2017) and prior study (Essen et al., 2013). In addition, we also obtained the group-average dense T1w/T2w myelination index provided by Rosen and Halgren (2021), which also originated from the HCP.

#### 2.1.2 Cortical parcellation and network assignment

Both structural and functional connectivity are based on the HCP multimodal parcellation scheme (HCP-MMP1.0), which consists of 180 cortical parcels per hemisphere (M. Glasser et al., 2016). In the study conducted by Rosen and Halgren (2021), this parcellation scheme was projected from the Workbench (Marcus et al., 2011) 32k grayordinate template to the FreeSurfer (Fischl, 2012) ico5 fsaverage template to perform tractography and calculate structural connectivity. In our work, we applied the same fsaverage space parcellation to calculate functional connectivity. Additionally, we utilized the network assignment provided by Rosen and Halgren (2021) for subsequent analysis. The network assignment which consists of 10 functional networks (Figure 1) was generated by simplifying the assignment scheme from Ji et al. (2019). Details on cortical parcellation and network assignment could be found in Rosen and Halgren (2021).

**Figure 1:**
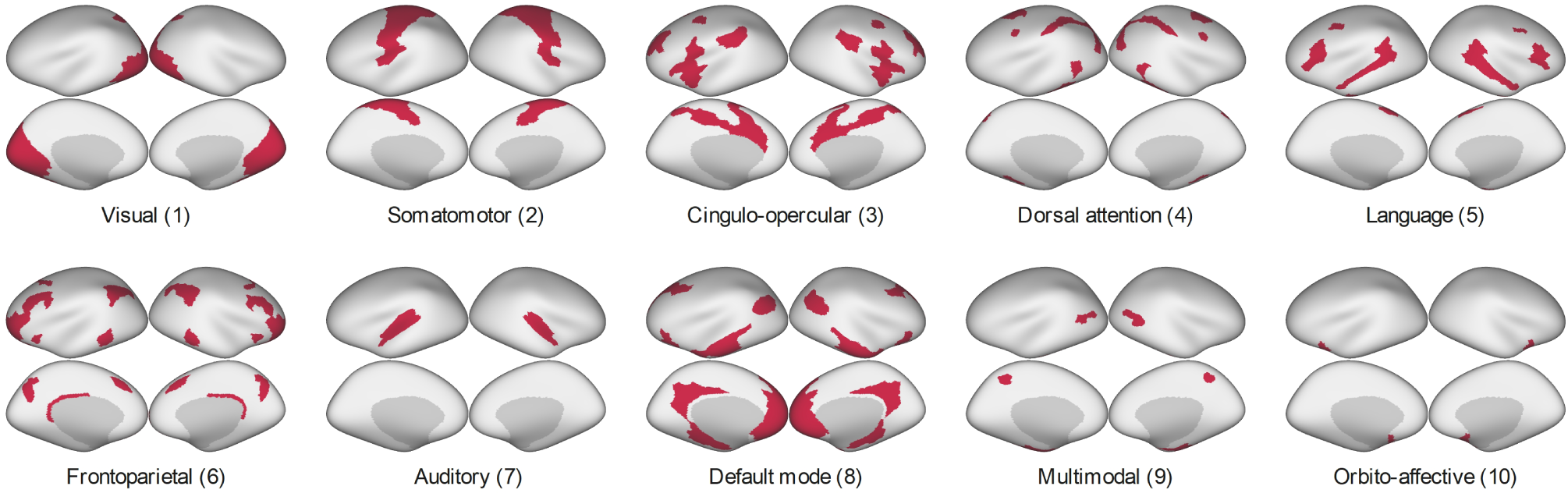
The network assignment scheme generated by Rosen and Halgren (2021). This scheme was based on the prior study (Ji et al., 2019). Specifically, the order and groupings of the left hemisphere in Ji et al. (2019) were used for homologous parcels in the both right and left hemisphere. Also, Rosen and Halgren (2021) simplified the network pairs by combining the two pairs of the original networks (primary and secondary visual, ventral, and posterior multimodal) into visual and multimodal groups.

#### 2.1.3 Structural connectivity

In the study conducted by Rosen and Halgren (2021), the diffusion MRI data were processed as follows to calculate structural connectivity (SC). First, the diffusion MRI data were corrected for eddy currents and movement with FSL eddy (Andersson & Sotiropoulos, 2016). Then fractional anisotropy (FA) analysis was performed using dtifit, and the resulting FA volumes were used to register the FreeSurfer and diffusion MRI volumes (flirt). Non-invasive probabilistic tractography was performed with probtrackx2 in voxel-byparcel mode (-os2t -s2tastext). The list of target parcels (-targetmasks) was quartered into four sublists, therefore, probtrackx2 was invoked 1440 times per subject, estimating the connectivity between 1 seed parcel and 90 target parcels in each invocation. The default 1/2 voxel step length, 5000 samples and 2000 steps were used (-step-length 0.5 -P 5000 -S 2000). To avoid artifactual loops, streamlines that loop back on themselves were discarded (-l) and tractography was constrained by a 90° threshold (-c 0) for maximal curvature between successive steps. More details on tractography can be found in the study (Rosen & Halgren, 2021).

For normalization, Rosen and Halgren (2021) applied fractional scaling to the structural connectivity generated by tractography. The fractionally scaled connection between two distinct regions was defined as the number of streamlines starting in A and ending in B, divided by the total number of streamlines connected to either A or B while excluding within-parcel connections. Subsequently, they symmetrized the structural connectivity matrix by taking the arithmetic mean of the A-to-B and B-to-A fractionally scaled connection weights. Additionally, the authors noted that probabilistic tractography values follow an approximately lognormal distribution, thus requiring log-transformation prior to subsequent analysis. However, the structural connectivity data obtained from Rosen and Halgren (2021) were not log-transformed. Therefore, in this study, we first log-transformed (log10) the obtained structural connectivity data, followed by Min-Max scaling to the range [0, 1], then we sparsified the structural connectivity by 50%, that is, we retained the first 50% of the strong connections to reduce computational overhead. Figure 2 illustrates the entire data preprocessing process.

**Figure 2:**
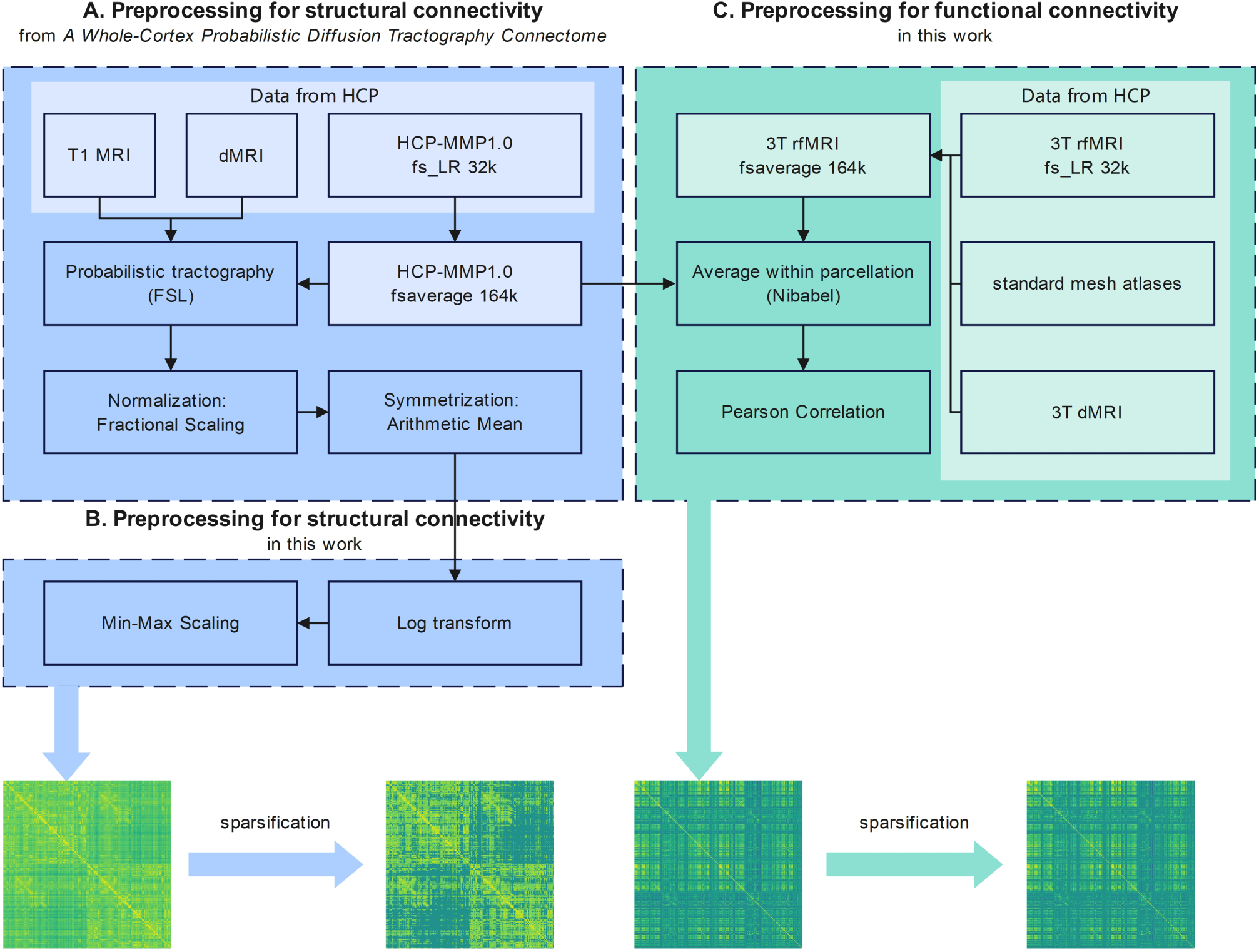
The entire data preprocessing. **A**, The preprocessing performed by Rosen and Halgren (2021), which includes tractography using structural MRI, diffusion MRI, and parcellation, followed by fractional scaling and symmetrization. **B** and **C**, The preprocessing for structural connectivity and functional connectivity in this work, which includes log-transformation and min-max scaling performed on the obtained structural connectivity, and resampling rs-fMRI from fs LR space to fsaverage space and calculating functional connectivity using the parcellation provided by Rosen and Halgren (2021). The light-colored areas represent data, and the dark-colored areas represent processing.

#### 2.1.4 Functional connectivity

To maintain consistency with structural connectivity, we utilized the 164k fsaverage HCP-MMP1.0 parcellation provided by Rosen and Halgren (2021), therefore, we first mapped individual rs-fMRI in the fs LR space to the fsaverage space according to (Coalson et al., 2017). This process was performed with Connectome Workbench version 2.1.0 (Marcus et al., 2011). We used the fsaverage-registered individual native sphere (-surface-sphere-project-unproject) to resample the individual’s native surface to 164k fsaverage mesh (-surface-resample), and then used this surface to map the separated 32k fs LR rs-fMRI (-cifti-separate) to 164k fsaverage (-metric-resample). Then we extracted the regional average BOLD signal based on the parcellation provided by Rosen and Halgren (2021). Lastly, functional connectivity (FC) was calculated as the Pearson correlation coefficient matrix between the BOLD time series of all 360 regions. As the resulting FC values were already approximately normally distributed, no further transformations were applied. Similar to SC, we also sparsified FC by 50%, retaining the top 50% of strong connections by absolute value. Figure 2 illustrates the entire data preprocessing process.

### 2.2 Approaches for calculating structure-function coupling

#### 2.2.1 Correlational approach

The correlational approach is the most commonly used approach for assessing structure-function coupling (SFC), in which the SFC of each cortical region was defined as the correlation coefficient (Pearson or Spearman) between its vectors of structural and functional connectivity (Baum et al., 2020; Gu et al., 2021; Honey et al., 2009; Liégeois et al., 2020). Generally, the connectivity vector can be the upper triangular element or row vector of the connectivity matrix, used to calculate the global SFC or the regional SFC, respectively. In this study, we used the Pearson correlation coefficient of the corresponding row vectors of the connectivity matrix to calculate the regional SFC, and in the subsequent analysis, the correlational approach was used as the baseline approach for the methodological difference, which refers to the difference produced without introducing actual indirect connections. The detailed definition of methodological difference will be explained later.

#### 2.2.2 Regression approach

In the study conducted by Vázquez-Rodríguez et al. (2019), multiple linear regression was used to calculate SFC. For each cortical region *i*, the dependent variable of regression model was the functional connectivity strength between region *i* and all other cortical regions *j* (*j* ≠ *i*), represented by the *i*^th^ row of FC matrix. The independent variables were the geometric and structural relationship metrics between regions *i* and *j*, including the Euclidean distance, shortest path length, and communicability. In this study, the communicability was defined as

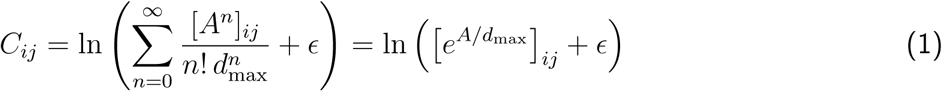

Where *A* is the binarized SC, and *d*_max_ is the maximum degree. Compared to the definition of communicability in Vázquez-Rodríguez et al. (2019), we normalized it by dividing the maximum degree and performed a log-transformation because the parcellation we used was larger (360 × 360), resulting in a larger graph size. Each *i* − *j* node pair constitutes an observation, therefore, the regression model for each node has 360 samples. The regression coefficients for each independent variable were estimated using ordinary least squares, and the model’s adjusted goodness of fit was calculated as the SFC for the corresponding cortical region. The coupling relationship between SC and FC is often highly nonlinear (Sporns, 2011), but when we introduce network characteristics, linear model can also be used to measure the correspondence between SC and FC to some extent. With proper design, their predictive ability can even be better than that of nonlinear models (Abdelnour et al., 2014). In this study, multiple linear regression was the simplest predictive model. Since the independent variables of the model come from the binarized SC, its predictive ability is actually weaker than other deep learning models. However, this also means that the informational difference arising from indirect connections only affect the predicted FC through topological structure. This is significantly different from the way other predictive models introduce informational difference, thus helping to find a more general informational difference pattern among predictive models.

#### 2.2.3 Multilayer perceptron

Multilayer perceptron (MLP) is a fundamental deep learning model that achieves global interaction between features through fully connected neurons. This characteristic makes it exhibit strong predictive ability when predicting FC. In this study, we used the MLP architecture from Sarwar et al. (2021), which contains four hidden layers, each containing two fully connected layers that use LeakyReLU and Tanh as activation functions respectively (each consists of 2048 neurons with a dropout rate of 0.5). The objective function was given by

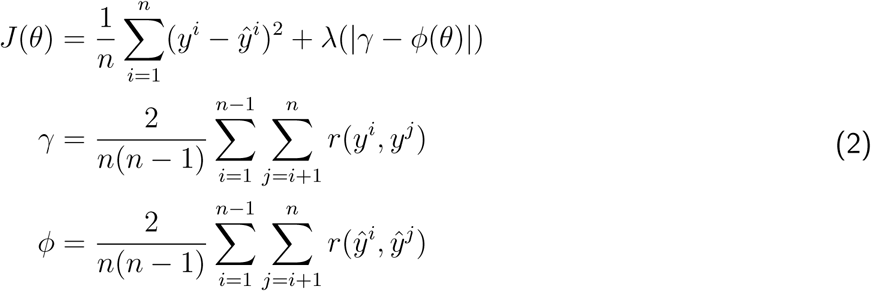

where *θ* denotes parameters of MLP, *i* indexes a subject. *y* is the empirical FC and *y*^ is the predicted FC. *r* represents the calculation of the Pearson correlation coefficient. The first term in the formula of the objective function is mean square error, and the second term is the regularization function which ensures that the MLP learned the mapping between SC and FC while preserving inter-individual differences. The upper triangular elements of SC were flattened and used as input to MLP to predict the upper triangular elements of FC. Then we calculated the Pearson correlation coefficient between the predicted FC and the empirical FC as SFC. In this study, our modifications to the model are mainly in two aspects: first, considering the computational burden, we simply doubled the number of neurons to accommodate higher-dimensional features resulting from more cortical regions (68 regions and 1024 neurons in Sarwar et al. (2021)); second, we added batch normalization layers to alleviate gradient vanishing.

We divided the 998 subjects who had both SC and FC into training set (898 subjects) and test set (100 subjects). 10-fold cross-validation was used to adjust the hyperparameters on the training set and finally all samples from the training set were used for training. Specifically, the model was initialized using the Kaiming Uniform and trained using the Adam optimizer (learning rate of 0.001 and weight decay of 10^-5^) for 4 epochs (batch size = 16). Notably, the number of samples used in training set and test set was actually four times the number of subjects because we did not merge the four runs of rs-fMRI, but kept them as samples. In this case, the SC of the same subject will be used to predict the FC of four different runs, which is actually equivalent to data augmentation and helps to improve the robustness of the model. The test set samples were used to calculate predicted FC and SFC for subsequent analysis, and finally were also used for individual identification. All deep learning models used in this study used the same training and test sets.

#### 2.2.4 Predictive GCN

In recent years, graph neural networks (GNNs) have been used in several studies to predict functional connectivity because they can extract features while preserving graph structure and integrate direct and indirect connections in brain networks (Chen et al., 2024; Neudorf et al., 2022; Wein et al., 2021). Graph convolutional network (GCN) is one of the most commonly used GNN, and predictive GCN (pGCN) refers to GCNs used in connection prediction tasks in this study. GCN performs feature extraction by applying aggregation and update operations on a graph without altering its structure (Kipf & Welling, 2017). It is computed as

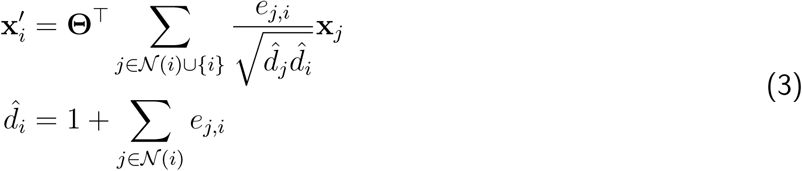

Here, **x**^′^ is the updated feature vector of node *i*, **Θ** denotes the trainable weights, N (*i*) represents the set of neighboring nodes of node *i* (excluding itself), *e_j,i_* is the edge weight from node *j* to node *i*, 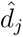 and 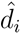 are the generalized degrees of node *j* and node *i*, respectively, i.e., the degrees after considering self-loops and edge weights. ∪{*i*} and 1 indicate the addition of self-loops. Self-loops allow nodes to consider their own features during aggregation.

As shown in the formula, a standard GCN is typically normalized by node degree. However, in this study, because both SC and FC were weighted graphs, we used the absolute weighted degree for normalization, which is

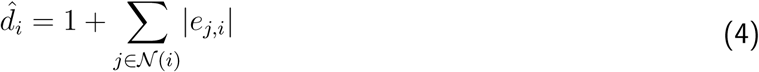

This allowed the model to be normalized correctly when FC was used as input in the self-supervised GCN and reverse predictive GCN, because there are negative connections in FC. During preprocessing, we set the diagonal elements of the SC matrix to zero because within-parcel connectivity was not examined (Rosen & Halgren, 2021), therefore, self-loops were not added for SC in GCN. In contrast, the diagonal elements of the FC matrix represent complete autocorrelation within each cortical region, thus, we retained the diagonal elements to add self-loops.

In this study, we reproduced the GNN architecture from Chen et al. (2024). In the model, a graph with structural connectivity strength as edge weights and one-hot vectors as node features was input into the GCN, and then the features of the connected endpoints were concatenated and fed into a MLP to predict the corresponding functional connectivity. Chen et al. (2024) used two methods to calculate individual SFC: one was to calculate the Pearson correlation between the upper triangular elements of the predicted FC and the upper triangular elements of the empirical FC as the whole-brain level SFC, and the other was to calculate the Pearson correlation coefficient between the *i*^th^ row of predicted FC and the empirical FC as the SFC of *i*^th^ region. Since the informational difference discussed in this study mainly stem from the introduction of indirect connections, we only calculated the second type of SFC (regional level) for analysis. The SFC calculated using other approaches also belong to this type of SFC.

Predictive GCN consists of a two-layer GCN (360 × 256 × 256) and a two-layer MLP (512 × 64 × 1). Mean squared error with L2-regularization was used as the loss function and Parametric Rectified Linear Unit (PReLU) was used as the activation function, which was defined by the following formula:

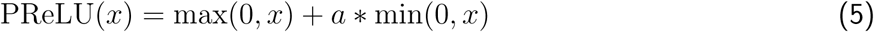

PReLU is a variant of the ReLU activation function, in which the coefficient *a* controls the slope in the negative region. The architecture of predictive GCN is consistent with that used by Chen et al. (2024), except that the dimension of the input features was modified to the number of cortical regions in the HCP-MMP1.0 atlas. Dropout (p = 0.5) and batch normalization layers were added to enhance regularization and prevent overfitting. For training, we used Kaiming Uniform initialization, and trained the model using Adam optimizer (learning rate of 0.001 and weight decay of 10^-5^) for 1 epoch (batch size = 16).

#### 2.2.5 Self-supervised GCN

To investigate the impact of simultaneously introducing indirect SC and indirect FC into the calculation of SFC, we designed self-supervised GCN (sGCN), which can demonstrate the additional effects of simultaneously introducing indirect SC and indirect FC in terms of the informational difference and can also be compared with the pGCN in terms of the methodological difference. Similar to the pGCN, connectivity matrix is represented as a graph with connection strengths as edge weights and one-hot vectors as initial node features, and then input into GCN. The difference is that SC and FC are simultaneously input into two GCNs with the same architecture, and then the feature vectors of SC nodes and FC nodes are output. The loss function of the sGCN is

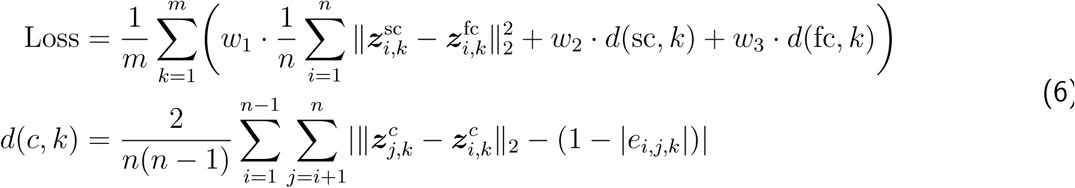

where *k* indexes a subject, *i* and *j* index nodes. ***z*** represents the feature vector output by the sGCN, *e_i,j,k_* is the edge weight between node *i* and node *j* of subject *k*. *c* indicates the modality to which the feature vector ***z*** belongs. *w_i_, i* ∈ {1, 2, 3} are hyperparameters that control the contribution of each loss component to the total loss. The first term of the loss function is the contrastive loss, and the second and third terms are the distance losses for SC and FC, respectively. Contrastive loss is used to establish a relationship between the feature vectors of corresponding nodes in SC and FC, while distance loss is used to constrain the feature space of the nodes. By minimizing this loss function, the distances between all corresponding SC nodes and FC nodes will be as close as possible, while the distances between nodes within the same modality will approach the actual connection strength. The SFC of cortical region *i* calculated using the sGCN was defined as the Gaussian kernel output of its feature vectors of SC and FC:

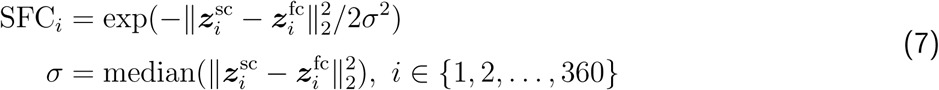

This definition is consistent with the optimization objective of sGCN, as both the contrastive and distance losses are calculated based on the Euclidean distance. The *σ* parameter for the Gaussian kernel was set as the median of the Euclidean distance between the SC and FC feature vectors across all cortical regions, ensuring robustness against potential outliers. Figure 3 illustrates the architecture of sGCN and how SFC is calculated. Both SC and FC parts of sGCN consist of a two-layer GCN (360 × 128 × 32) and were initialized using Kaiming Uniform. We added dropout (p = 0.2) and batch normalization layers for regularization. The weights for the loss components were set as: 1 for contrastive loss, 10 for SC distance loss, 15 for FC distance loss. Both SC and FC models were optimized using the Adam optimizer with identical hyperparameters (learning rate of 0.01 and weight decay of 0.001) for 1 epoch.

**Figure 3:**
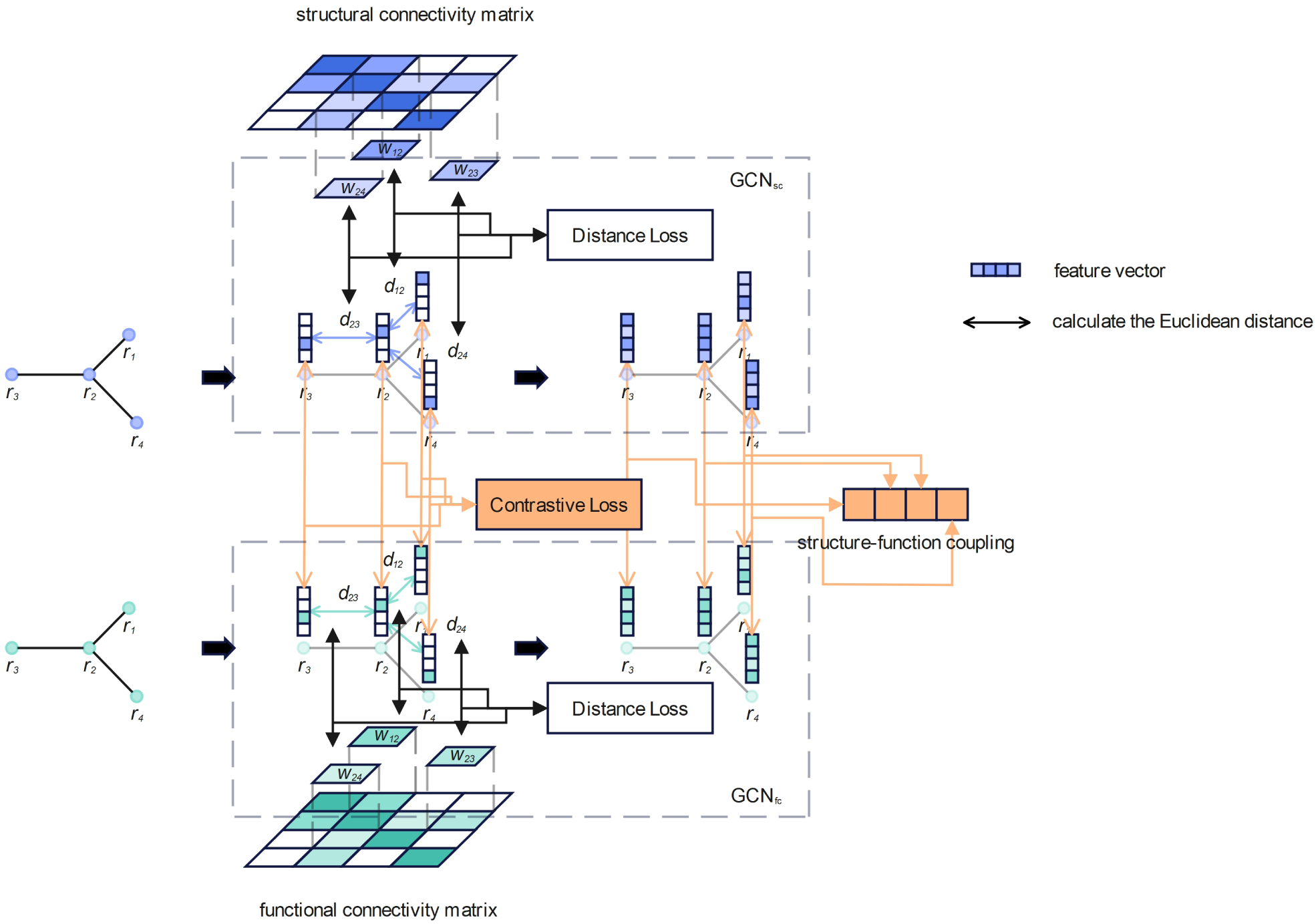
The architecture of self-supervised GCN (sGCN). This figure illustrates how sGCN uses the SC and FC with four cortical regions to calculate SFC. *r* represents the cortical region shown as a node in the graph. *w* is edge weight, i.e., the connection strength in SC or FC. *d* is the Euclidean distance between feature vectors of the nodes. sGCN adjusts parameters through training to continuously reduce contrastive loss and distance loss, ultimately making the distance between strongly connected cortical regions in the feature space closer, and making all corresponding SC and FC cortical regions as close as possible. In this example, the SFCs of four cortical regions are equal, because the SC and FC are exactly the same.

### 2.3 Comparison and analysis for informational difference and methodological difference

#### 2.3.1 Informational difference and methodological difference

In this study, we assume that for a given cortical region, the information involved in calculating the regional SFC only includes four types: direct structural connections, indirect structural connections, direct functional connections, and indirect functional connections. The informational difference we discuss refers to the difference arising from the use of different combinations of information by different approaches in calculating regional SFC. For example, the correlational approach for calculating the regional SFC using the row vectors of the connectivity matrix only uses two types of information: direct structural connectivity (SC) and direct functional connectivity (FC) (Baum et al., 2020). In contrast, the predictive model using the correlation coefficient between the corresponding row vectors of the predicted FC and the empirical FC as regional SFC utilizes three types of information: direct SC, indirect SC, and direct FC (Chen et al., 2024; Vázquez-Rodríguez et al., 2019). It needs to be clarified that some deep learning models do capture indirect FC when predicting FC (Chen et al., 2024; Neudorf et al., 2022; Sarwar et al., 2021), but this is because the indirect FC of the training samples affect the model’s weights, rather than taking the actual indirect FC into account during prediction. Therefore, in this study, the differences introduced through training samples are categorized as methodological differences, not as informational differences. In summary, the informational difference mentioned in this study reflects whether actual direct or indirect connections are used when calculating SFC.

Of the five approaches used in this study to calculate SFC, three are categorized as predictive approaches (Figure 4 A). In these three predictive approaches, the regional SFC of the regression approach was defined as the adjusted goodness of fit between the predicted FC and the empirical FC, while the regional SFC of the MLP and the pGCN was defined as the Pearson correlation coefficient between the predicted FC and the empirical FC. In these three predictive approaches, the regional SFC of each cortical region is calculated from its SC vectors and FC vectors, which means that only direct connections of the predicted FC and the empirical FC are involved in the SFC calculation. In predictive approaches, the calculation of the predicted FC involves both direct and indirect SC. Therefore, in general, the information involved in calculating the regional SFC in these three predictive approaches includes direct SC, indirect SC, and direct FC. If the predictive model is run in reverse, i.e., using FC to predict SC, the information involved in the regional SFC calculation includes direct FC, indirect FC, and direct SC. In the remaining two non-predictive approaches (Figure 4 B), the correlational approach only uses direct SC and direct FC to calculate the regional SFC. The SFC calculated using the sGCN was defined as the Gaussian kernel output of the structural vector representation and the functional vector representation. Since the calculation of these two vector representations involves direct and indirect connections, the sGCN uses all the information when calculating the regional SFC. Table 1 shows the information combinations used in the approaches to calculating SFC in this study.

**Table 1:** The information combinations used in each approach

| Approach | Information types |  |  |  |
| --- | --- | --- | --- | --- |
|  | direct SC | indirect SC | direct FC | indirect FC |
| Correlation / direct predictive models<sup>a</sup> / direct sGCN | ✓ |  | ✓ |  |
| predictive models / full-SC sGCN | ✓ | ✓ | ✓ |  |
| reverse<sup>b</sup> predictive models / full-FC sGCN | ✓ |  | ✓ | ✓ |
| sGCN | ✓ | ✓ | ✓ | ✓ |

<sup>a</sup> predictive models: Regression/MLP/pGCN
<sup>b</sup> reverse: refers to using FC to predict SC.

The methodological difference refers to the difference that persists even when calculating SFC using the same information combination in different approaches. However, since many approaches actually use the same information combination (table 1), we used the SFC calculated using the correlational approach as a unified baseline to calculate the methodological difference.

#### 2.3.2 Full model and direct model

Due to the different properties of direct and indirect connections, many studies have attempted to illustrate their different roles in predicting FC by considering only direct or indirect connections (Esfahlani et al., 2022; Røge et al., 2017). In this study, to exclude the methodological difference when calculating the informational difference and to compare each approach with the correlational approach, we also attempted to calculate SFC using only direct connections in each approach. Specifically, for each cortical region, we retrained the models (regression, MLP, pGCN, and sGCN) using samples with indirect connections excluded, thereby calculating the predicted connectivity and SFC that only considered direct connections. In Figure 4, the direct pFC and the direct SFC refer to the predicted FC and SFC calculated by the direct models, while the full pFC and SFC refer to the predicted FC and SFC calculated by the original models that retained indirect connections (full models). It should be noted that since a complete sGCN uses all direct and indirect connections, there are also sGCNs trained using the samples with only indirect FC or indirect SC connections excluded (full-SC sGCN and full-FC sGCN).

When training a direct model for the *i*^th^ cortical region, we retained the *i*^th^ row and the *i*^th^ column of its connectivity matrix while setting other elements to zero to exclude indirect connections. However, for the MLP, excluding indirect connections in this way leads to very sparse SC and FC vectors, resulting in a large number of redundant parameters. Furthermore, for the MLP, simply setting the indirect connections to zero does not mean completely ignoring them. Therefore, when training direct MLP for the *i*^th^ cortical region, we used the *i*^th^ row of the connectivity matrix as input to predict the *i*^th^ row of the connectivity matrix of the other modality, while adjusting the number of neurons in the hidden layers to 128 to accommodate the lower-dimensional input. The other designs and hyperparameters of the MLP remain unchanged.

#### 2.3.3 Differences at the predicted-connectivity level

In this study, we attempted to analyze how the informational and methodological differences are manifested at the predicted-connectivity level and the SFC level (Figure 4 C). The first level was set for the predictive approaches. Taking the direction of using SC to predict FC as an example, the most significant characteristic of the predictive approaches is that the predicted FC is used to replace SC in calculating the correlation coefficient with the empirical FC (Chen et al., 2024; Sarwar et al., 2021; Vázquez-Rodríguez et al., 2019). This means that the informational and methodological differences are primarily introduced from the predicted FC. Therefore, compared to the SFC level, the differences at the predicted-connectivity level are closer to the source of the differences and can more precisely reflect the impact of these two types of differences.

The informational difference and the methodological difference at the predicted-connectivity level were calculated using the following formulas.

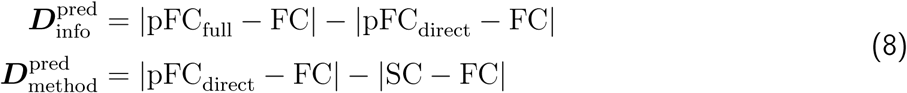

pFC_full_ and pFC_direct_ are the predicted FC calculated by the full models and the direct models, respectively, as mentioned in Section 2.3.2. For predictive models with the direction of using FC to predict SC, the differences were calculated in the same way, simply by interchanging the SC and FC modalities. The formulas defined the informational difference as the change in the absolute error of the prediction after introducing indirect connections, while the methodological difference was the change in the absolute error with only direct connections considered relative to the original error between SC and FC. Because each connection interacts with different connections when acting as a direct connection of different cortical regions, neither pFC_direct_ nor pSC_direct_ is symmetric. To ensure that the differences in each connection are correctly estimated, we symmetrized pFC_direct_ and pSC_direct_ by taking the arithmetic mean of symmetric connections when calculating the differences at the predicted-connectivity level. However, we retained the asymmetric pFC_direct_ and pSC_direct_ when subsequently calculating the SFC.

#### 2.3.4 Differences at structure-function coupling level

At the SFC level, we took the difference between the SFC calculated by the full model and the direct model as the informational difference, and the difference between the SFC calculated by the direct model and the correlational approach as the methodological difference. Therefore, the differences were defined as:

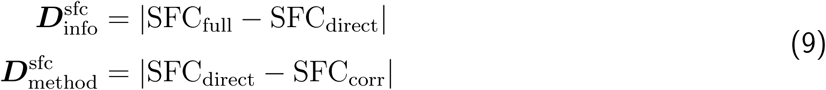

For predictive approaches, the differences of the models with the direction of using FC to predict SC were also calculated in the same way. Similarly to the two prediction directions of the predictive approaches (SC-to-FC and FC-to-SC), the informational difference of the sGCN also includes two types: |SFC_full-sc_ − SFC_direct_| and |SFC_full-fc_ −SFC_direct_|. SFC_full-sc_ and SFC_full-fc_ represent the SFC calculated using full-SC sGCN and full-FC sGCN (defined in Section 2.3.2), respectively. All of these informational differences reflect the changes in SFC after the first introduction of indirect connections, i.e., the changes in SFC calculated by the approaches in the second and third rows of Table 1 relative to the SFC calculated by the approaches in the first row. Furthermore, for the sGCN, we also calculated |SFC_full_ − SFC_full-sc_| and |SFC_full_ − SFC_full-fc_| to illustrate how the second introduction of indirect connections affects SFC.

To visually demonstrate the relationship between SFC calculated using different approaches, we performed dimensionality reduction using principal component analysis (PCA) (Jolliffe, 2014) on the SFC calculated by each approach. To further compare the informational difference and the methodological difference, and to ensure statistical significance of our conclusions, we performed two-sided paired t-tests on the differences for each region, and the Benjamini–Hochberg procedure (Benjamini & Hochberg, 1995) was used to control the false discovery rate (FDR) at 5% (i.e., q *<* 0.01).

#### 2.3.5 Individual identification

To investigate the impact of the informational difference and the methodological difference on identification accuracy when using SFC to distinguish individuals, we used the SFC calculated by different approaches (including direct models) for individual identification. Specifically, for the SFC of the test set (100 subjects, 4 runs per subject) calculated by each approach, we iterated the number of subjects *n* from 10 to 100 (with a step size of 1). For each *n* value, we randomly selected *n* subjects from the 100 subjects to form a subset. For each subset, we used support vector classifier (SVC) (Cortes & Vapnik, 1995), random forest (RF, n estimators = 100) (Breiman, 2001), and multilayer perceptron (MLP, 128 × 64 × 32) to classify SFC, and evaluated the classification accuracy using 20 repeated 3-fold cross-validation. Finally, we reported the overall distribution of the accuracy of the SFC calculated by each approach in these 91 individual identifications with different numbers of subjects.

## 3 Results

In this study, we obtained ICA-FIX denoised rs-fMRI data comprising 1003 subjects (469 males, 22-36+ years old) from the WU-Minn Human Connectome Project 1200 subjects release (HCP S1200) (Essen et al., 2013) and structural connectivity (SC) data comprising 1065 subjects (490 males, 22-36+ years old) of HCP S1200 from Rosen and Halgren (2021). To be consistent with SC, we mapped the rs-fMRI data from the fs LR space to the fsaverage space and calculated the Pearson correlation coefficient matrix between the regional average time series as functional connectivity (FC). The calculation of SC and FC was based on the HCP multimodal parcellation scheme (HCP-MMP1.0, 180 regions per hemisphere) (M. Glasser et al., 2016), so the SC and FC of each subject are 360 × 360 symmetrical matrices. For the obtained SC, we performed log-transformation to ensure the normality and Min-Max scaling for normalization. Both SC and FC were sparsified by 50%, which means that only the top 50% of connections with the highest strength were retained. Finally, 898 of the 998 subjects who possessed both SC and FC were used as training set, while the remaining 100 subjects served as test set.

We used five approaches to evaluate structure-function coupling (SFC), including three predictive approaches (Figure 4 A): regression approach (Vázquez-Rodríguez et al., 2019), multilayer perceptron (MLP) (Sarwar et al., 2021), and predictive graph convolutional network (pGCN) (Chen et al., 2024), and two non-predictive approaches (Figure 4 B): correlational approach (Baum et al., 2020; Gu et al., 2021; Honey et al., 2009; Liégeois et al., 2020) and self-supervised graph convolutional network (sGCN). Detailed definitions of these five approaches are given in Section 2.2. We then explained that these five approaches use different combinations of information (Table 1) when calculating regional SFC and how to exclude indirect connections to acquire predicted connectivity and SFC with only direct connections considered (direct model).

To systematically analyze the differences between different approaches in calculating SFC, we first defined the scope of the informational difference and the methodological difference. Based on the division of information combinations (Table 1), we described the informational difference as the difference that arises when calculating the regional SFC due to the use of different combinations of information by different approaches. And the methodological difference refers to the difference that still exists even when the same combination of information is used in the calculation of regional SFC. Then we calculated the specific informational and methodological differences at the predicted-connectivity level and SFC level (Figure 4 C).

It is worth noting that we calculated these differences for both two prediction directions (SC-to-FC and FC-to-SC) of the predictive models. Furthermore, to further illustrate the impact of these differences, we performed individual identification using SFC calculated by different approaches and presented the overall distribution of identification accuracy for each approach across different numbers of subjects.

### 3.1 Modeling approaches capture effective structure-function coupling

The study conducted by Vázquez-Rodríguez et al. (2019) using multiple linear regression has shown that the primary sensory and motor cortices exhibit high structure-function correspondence, while the transmodal cortex exhibits low correspondence. In this study, the structure-function correspondence of the visual network calculated using our reproduced multiple linear regression model was the highest, while the correspondences of the auditory, cingulo-opercular and language networks were the lowest, which is consistent with the results given by Vázquez-Rodríguez et al. (2019) using the same model (Figure 5 B and Supplementary Figure S1 B). The multilayer perceptron (MLP) we reproduced from Sarwar et al. (2021) also exhibits similar results, with higher structure-function coupling (SFC) in the visual and somatomotor networks, and lower in the orbito-affective, language, and auditory networks (Figure 5 B and Supplementary Figure S1 C). As reported by Sarwar et al. (2021), the MLP can achieve superior performance in predicting FC. Consistently, in our reproduction of their approach, the predicted connectivity was highly similar to empirical connectivity (individual: *r̅* = 0.69 ± 0.1), whether predicting FC from SC or predicting SC from FC (Figure 5 C and Figure 5 D).

**Figure 4:**
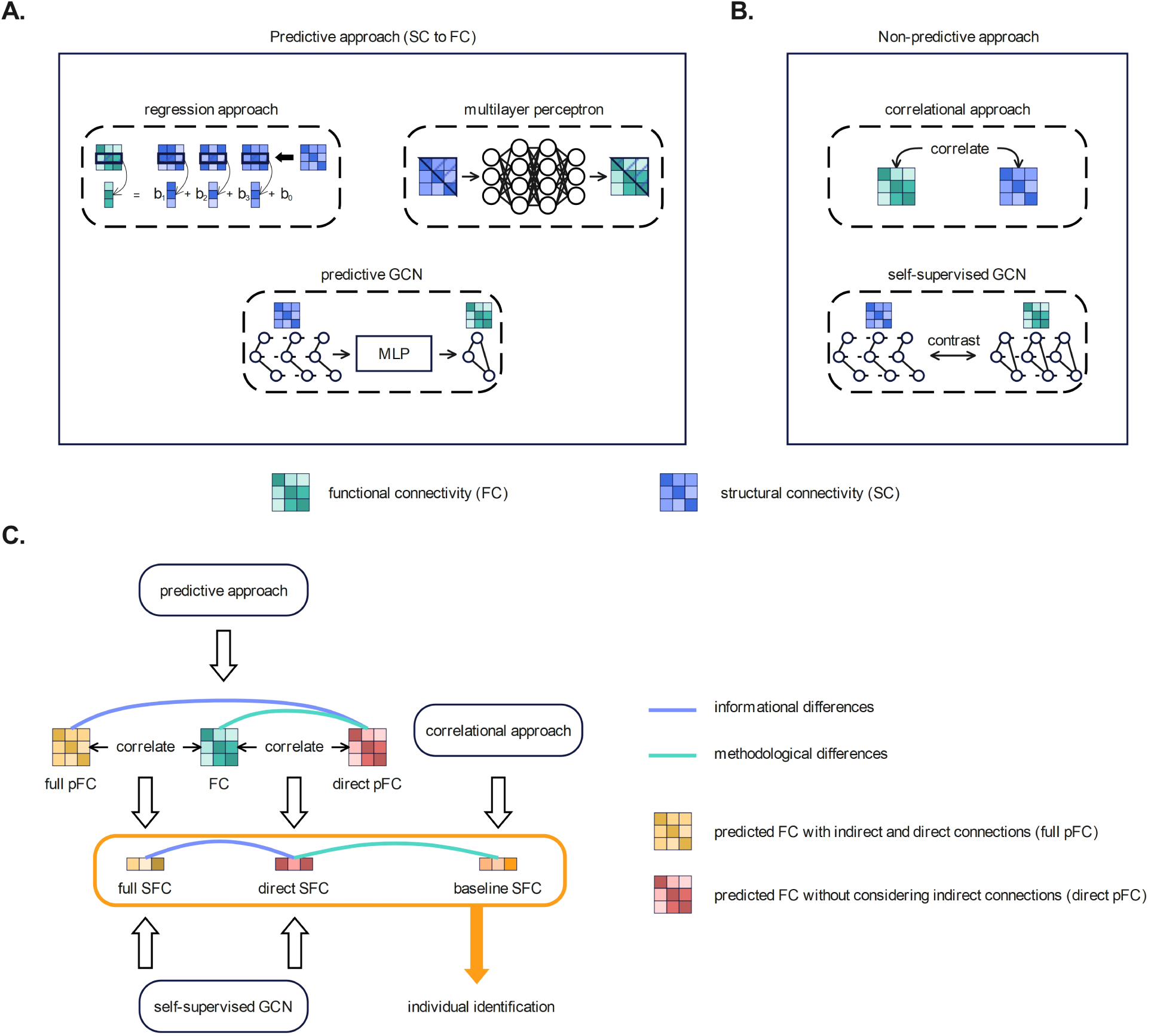
Schema of study design and methodology. **A** and **B**, Five approaches used to calculate SFC in this study. For a detailed definition of each approach, see Section 2.2. **C**, The process of analyzing informational differences and methodological differences. The definitions of the two differences are given in Section 2.3.1. In this study, we explore the informational and methodological differences at two levels: predicted connectivity (for predictive models) and structure-function coupling (SFC). Sections 2.3.3 and 2.3.4 respectively explain how the differences within each of the two levels are calculated. Finally, the SFC calculated by each approach was used for individual identification to explore the impact of informational and methodological differences on individual identification tasks. This schematic only considers one direction of the predictive approaches (using SC to predict FC), but in the actual analysis, we also calculated the two differences in the other direction (using FC to predict SC).

**Figure 5:**
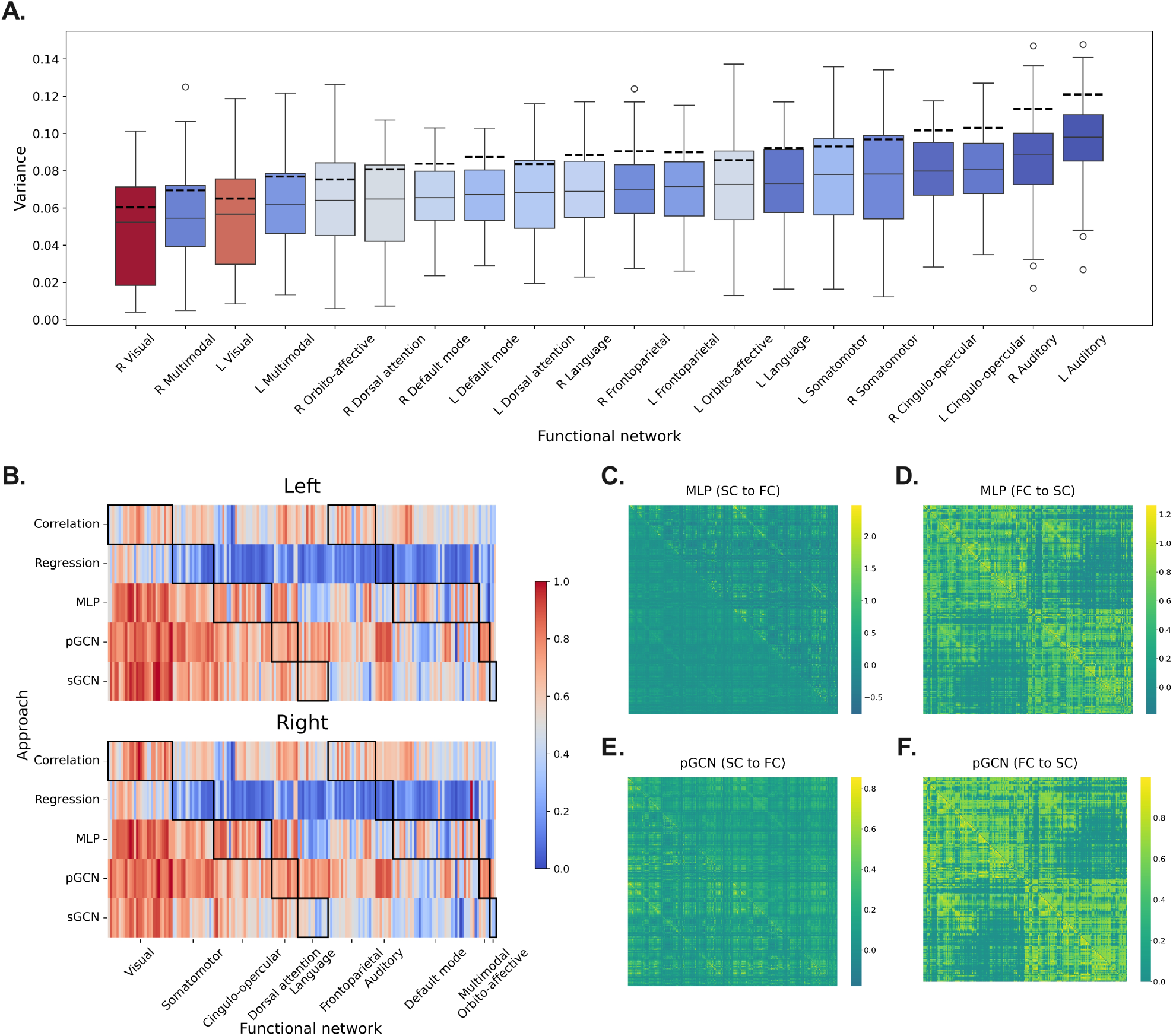
Effectiveness of the five approaches (see Section 2.2 for a detailed definition of each approach). **A**, Relationship between absolute deviation and cross-approach consistency in network-level structure-function coupling (SFC). Each box represents the distribution, over all test samples, of the variance among the five SFC values calculated from the different approaches for each functional network. The color of each box represents the absolute deviation of each network’s mean SFC from the overall mean SFC across all networks; the closer the color is to red, the larger the absolute deviation, and the closer to blue, the smaller the absolute deviation. The dashed line of each box represents the median of the variance distribution of the four remaining approaches after removing the sGCN. SFC were scaled using Min-Max scaling before calculating cross-approach variance and mean. **B**, The regional SFC after Min-Max scaling calculated using the five approaches (see Supplementary Figure S1 for cortical map of SFC). Neither **A** nor **B** involves the predictive models with the direction of predicting SC from FC. **C**, **D**, **E**, and **F**, Comparison between predicted connectivity and empirical connectivity. In SC-to-FC prediction, the lower triangle of the matrix represents empirical FC, while the upper triangle represents predicted FC. Conversely, for FC-to-SC prediction, the lower triangle corresponds to empirical SC, and the upper triangle corresponds to predicted SC.

Among the three predictive models used in this study, the predictive GCN (pGCN) achieved the highest prediction accuracy at the individual level (*r̅* = 0.75 ± 0.2), effectively capturing the structure-function mapping relationship (Figure 5 E and Figure 5 F). Furthermore, consistent with the results of Chen et al. (2024), in our reproduction, the visual, somatomotor, and auditory networks exhibited high SFC (with the visual network being the highest), while the frontoparietal and default mode networks showed lower SFC (Figure 5 B and Supplementary Figure S1 D). Similar to other approaches, our self-supervised GCN (sGCN) calculated higher SFC for the visual network, while lower SFC for the default mode and frontoparietal networks (Figure 5 B and Supplementary Figure S1 E). Moreover, the presence of the sGCN reduces the cross-approach variance for all networks (Figure 5 A), indicating that the SFC calculated by the sGCN are more consistent with the central tendency of the SFC calculated by the other four approaches.

Previous studies have shown that T1w/T2w ratio gradually decreases along the sensorimotor-association axis from the primary sensorimotor cortex to the association cortex (Baum et al., 2022; M. F. Glasser & Essen, 2011), and SFC also decreases along this axis (Collins et al., 2024; Vázquez-Rodríguez et al., 2019). Therefore, we also calculated the Pearson correlation coefficient between the SFC calculated by each approach and T1w/T2w myelination index. The results show that all full models that predict FC from SC, as well as the SFC calculated by the correlational approach and the (full) sGCN, are positively correlated with myelination index (Table 2), which confirms the view that SFC decreases along the sensorimotor-association axis. In addition, we also observed that the direct and full models of the regression approach and the MLP did not differ much in their correlation with myelination index, but the difference was large in the pGCN and the sGCN, suggesting that the introduction of indirect connections has a greater influence in the GCN models. Subsequent analysis of the differences at SFC level will further prove this point.

**Table 2:**
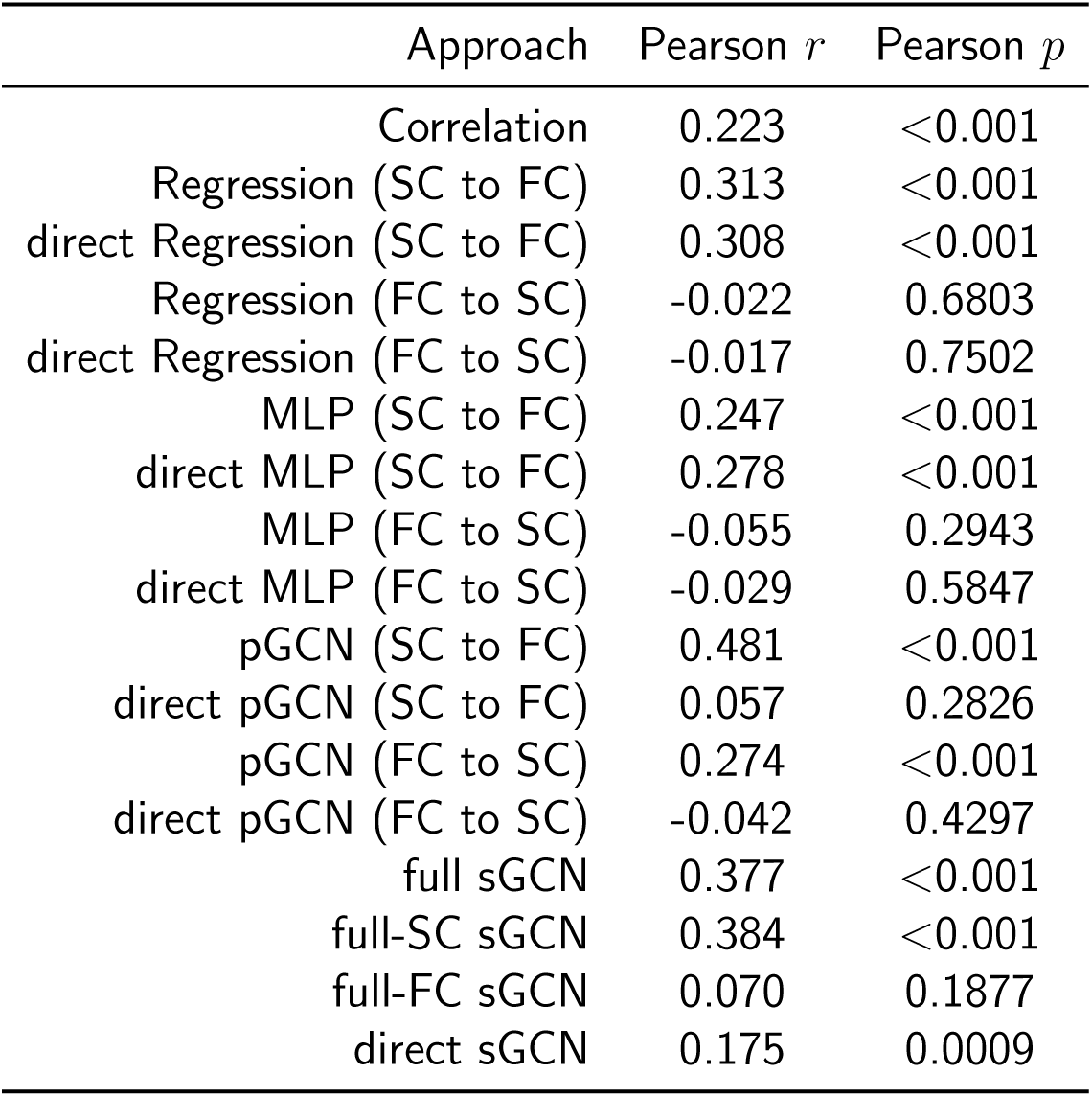
Pearson correlation coefficient between structure-function coupling and myelination index

The above description shows that modeling approaches all capture effective structure-function coupling patterns and exhibit a certain degree of consistency. Figure 5 A illustrates the relationship between absolute deviation and cross-approach consistency. All functional networks were arranged in ascending order of cross-approach variance. The color of each box represents the absolute deviation of each network’s mean SFC from the overall mean SFC across all networks, and the closer the color is to red, the larger the absolute deviation, i.e., the more extreme the SFC value. Figure 5 A indicates that, in general, networks with clearer structure-function relationships or more extreme SFC values, such as visual, orbito-affective, and default mode, exhibit higher cross-approach consistency. This observation aligns with common understanding, as clearer structure-function coupling is easier to capture.

Even though there is consistency in the SFC calculated by different approaches, the overall differences are significant, and the Pearson correlation coefficients among the SFC calculated by the five approaches are all below 0.6 (Supplementary Figure S1 F). As shown in Figure 5 B, significant differences in SFC calculated by different approaches still exist in some networks, such as the somatomotor, cingulo-opercular, and auditory. This suggests that different approaches may capture structure-function coupling patterns from different angles.

### 3.2 Differences at the predicted-connectivity level

After calculating the differences at the predicted-connectivity level, we performed preliminary statistical analysis on the results (Supplementary Figure S2). The results showed that most of the differences were negative and the proportion of negative values in the methodological differences was greater than that in the informational differences. This means that both the informational difference and the methodological difference contribute to predicting one modality from the other, but the reduction in prediction error mainly stems from the methodological difference, i.e., the properties of the model itself.

Furthermore, there were obvious differences between the differences of different models. Figure 6 (panels A-F) shows the mean difference matrices of the three predictive models on the test set. The lower triangle of each difference matrix represents the difference in predicting FC from SC, and the upper triangle represents the difference in predicting SC from FC. As can be seen in panels D and E, the methodological difference matrices of the regression approach and the MLP are quite similar, and both have better predictive performance for intra-hemispheric connections. Compared to the methodological difference, the informational difference matrices of the regression approach and the MLP have more positive values (Supplementary Figure S2 A, Figure 6 A and B), indicating that the introduction of indirect connections increases the prediction error of a considerable portion of the connections. This is particularly evident in the inter-hemispheric region of the upper triangle of panel A and in the contralateral homologs (secondary diagonal) and intra-network (diagonal) connections of the lower triangle of panel B. Overall, however, the impact of the informational differences on the regression approach and the MLP are relatively small (Supplementary Figure S2 B), regardless of whether the informational differences are positive or negative. In contrast, the informational difference plays a significant role in improving the predictive ability of the pGCN. Both the mean negative informational difference and the difference between the proportions of negative and positive values are obviously higher in the pGCN than in the other two models (Supplementary Figure S2 A and B). This suggests that indirect connections play a more important role in the pGCN.

**Figure 6:**
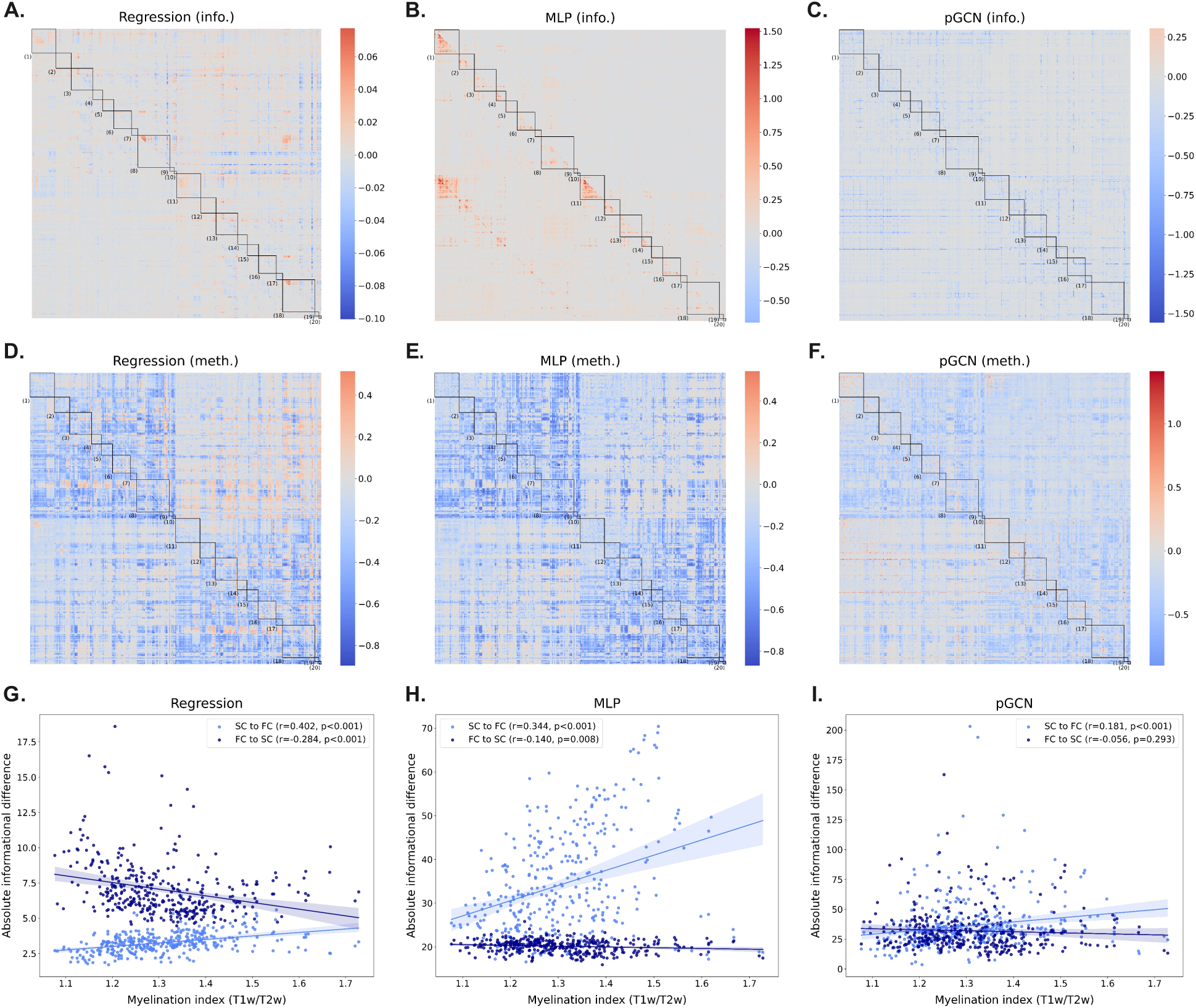
The difference matrices at the predicted-connectivity level and their correlation with myelination index. Panels **A**-**F**, The informational difference matrices and the methodological difference matrices of the three predictive models. The lower triangle of each difference matrix represents the difference in predicting FC from SC, while the upper triangle represents the difference in predicting SC from FC. The numbers in each matrix represent the corresponding functional networks (Figure 1). Panels **G**-**I**, The correlation between the sum of the absolute informational differences in direct connections of each cortical region and its myelination index.

To explore the relationship between differences and myelination index, for each cortical region, we summed the absolute informational differences in all direct connections and then calculated the means of the sums of differences on the test samples. Finally, the correlation coefficients between the total differences and myelination indices were calculated. The results showed that the informational difference of each predictive model was positively correlated with myelination index when predicting FC (Figure 6 G, H, and I). This suggests that the introduction of indirect connections has a greater impact on the prediction of direct connections around the cortical regions with higher myelination indices. In contrast, the methodological differences and the informational differences generated in SC prediction did not show a stable correlation with myelination index.

### 3.3 Differences at structure-function coupling level

After calculating the informational and methodological differences at SFC level, we found that the informational differences of the regression approach and the MLP were lower than their methodological differences (Supplementary Figure S3 A-H). In contrast, the pGCN and the sGCN exhibited both high informational and methodological differences (Supplementary Figure S3 I-Q). Overall, however, the four modeling approaches showed good consistency in cortical regions with low informational differences, and diverged in regions with high informational differences (Figure 7 C). After applying the functional network assignment provided by Rosen and Halgren (2021), we found that the orbito-affective network had the highest mean informational difference in both SC-to-FC and FC-to-SC directions, while the right-hemispheric dorsal attention network had the lowest (Figure 7 A). However, for most functional networks, their mean informational differences varied across different prediction directions and hemispheres. The former may stem from the different topologies of SC and FC, thus reflected in the informational differences of each of the four models, while the latter is mainly reflected in the GCN models, which may be related to the way network information is used.

**Figure 7:**
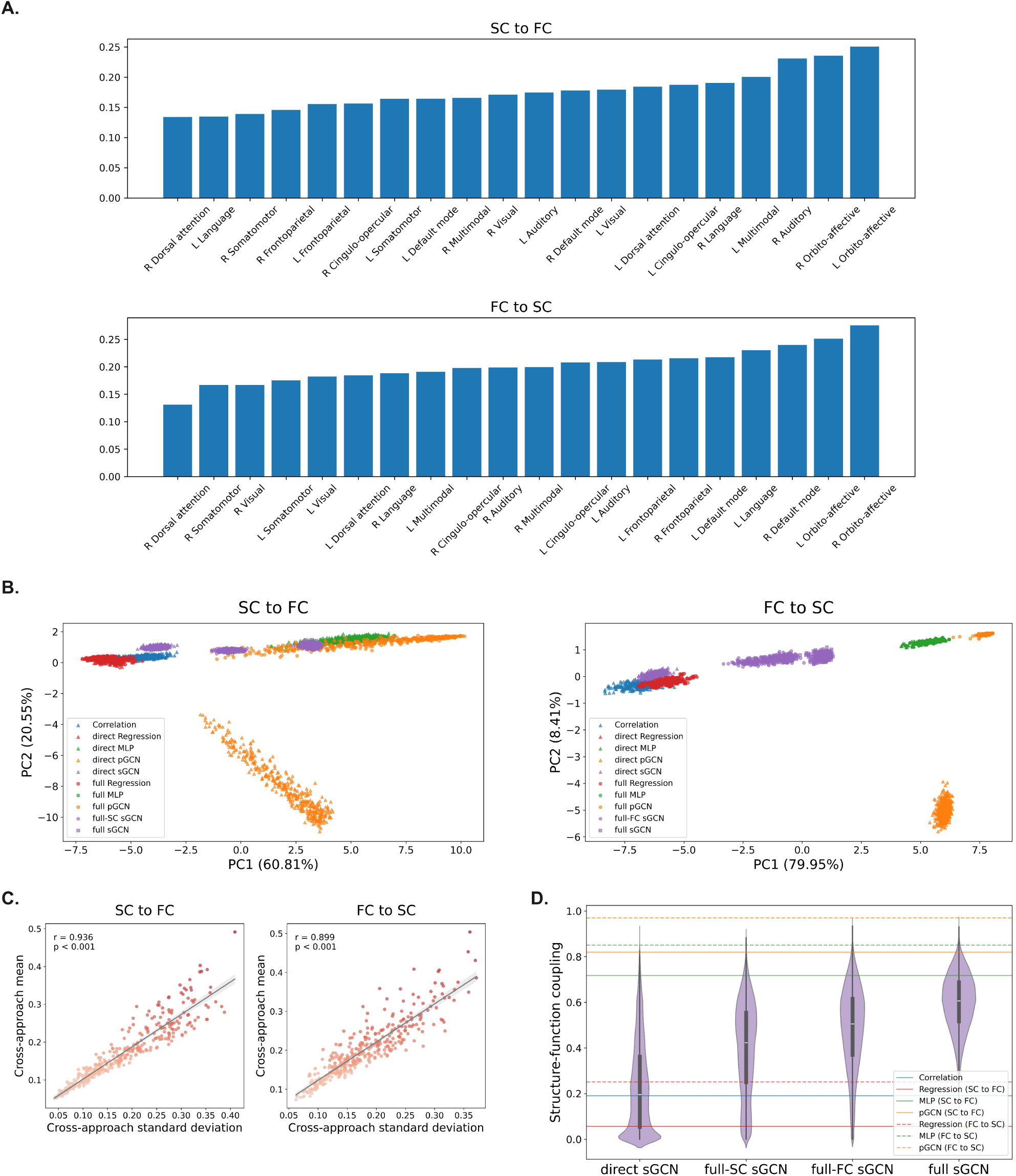
The informational difference at SFC level. **A**, the cross-approach mean of informational difference of functional networks at SFC level. **B**, Score plots of the first two principal components obtained by dimensionality reduction using principal component analysis (PCA) on SFC calculated by different approaches. **C**, Relationships between cross-approach mean and standard deviation of the informational differences. Panel **D** shows the distributions of SFC calculated by the sGCN with different information combinations.

To further quantify the relative magnitude of the informational difference and the methodological difference for each approach, we performed two-sided paired t-tests on the differences for each cortical region to determine the proportions of regions where the methodological difference was significantly higher than the informational difference and vice versa. The results are shown in Table 3. As shown in the table, for the regression approach and the MLP, the methodological differences are higher than the informational differences in almost all cortical regions, while in the GCN models, the cortical regions where the informational differences are higher than the methodological differences are obviously more numerous than those of the regression approach and the MLP.

**Table 3:** Comparison of methodological and informational differences across cortical regions: results of paired t-tests

| Approach | Meth. > Info. (%) | Meth. < Info. (%) | Significant regions<sup>a</sup>(%) |
| --- | --- | --- | --- |
| Regression (SC to FC) | 100.0 | 0.0 | 100.0 |
| Regression (FC to SC) | 100.0 | 0.0 | 100.0 |
| MLP (SC to FC) | 98.3 | 1.4 | 99.7 |
| MLP (FC to SC) | 100.0 | 0.0 | 100.0 |
| pGCN (SC to FC) | 75.3 | 23.3 | 98.6 |
| pGCN (FC to SC) | 92.2 | 6.7 | 98.9 |
| sGCN (full SC) | 41.4 | 52.5 | 93.9 |
| sGCN (full FC) | 26.4 | 66.7 | 93.1 |

<sup>a</sup> Significant regions: the proportion of cortical regions that remained significant after controlling the false discovery rate (FDR).

To more intuitively observe the relationship between the SFCs calculated by different approaches, we performed dimensionality reduction on them using principal component analysis (PCA) (Jolliffe, 2014). Figure 7 (panel B) shows the score plots of the first two principal components (PC1 and PC2). PC1 and PC2 cumulatively explained 81.4% and 88.4% of the total variance in the two prediction directions, respectively, which indicates that the first two principal components are sufficient to represent the overall structure of the original data. The results of PCA show that, for the regression approach and the MLP, the SFC calculated by the full model and the direct model are very similar in the first two principal components, indicating that the informational differences are very low, which is consistent with the t-test results. However, for the pGCN and the sGCN, the SFC calculated by the full model and the direct model differs significantly, forming independent clusters in the first two principal components. Furthermore, it can be observed that the SFC calculated by the direct sGCN is closer to that of the correlational approach than that of the full-SC sGCN and the full-FC sGCN, reflecting the characteristic that its informational difference outweighs the methodological difference, which is also reflected in the results of the t-tests. In addition, the PCA results indicate that the SFCs of the regression and the correlational approaches are relatively close, as are those of the MLP and the full pGCN, but along the second principal component, the SFC of the direct pGCN differs markedly from the others.

To investigate the impact of introducing indirect connections a second time in the calculation of SFC, we designed the sGCN model. The SFC calculated by the sGCN using different information combinations are shown in Figure 7 (panel D). The change from the direct sGCN to the full-SC sGCN in the figure corresponds to the first introduction of indirect connections in the SC modality, and also corresponds to the predictive model in the SC-to-FC prediction direction. The change from the full-SC sGCN to the full sGCN corresponds to the second introduction of indirect connections (FC modality) after the initial introduction in the SC modality. The same logic applies to the full-FC sGCN, the change from the direct sGCN to the full-FC sGCN and then to the full sGCN corresponds to the sequential introduction of indirect connections in the FC and SC modalities. We found that after introducing indirect connections, the SFC calculated by the sGCN gradually shifts from the correlational approach and the regression approach side (lower SFC) to the MLP and the pGCN side (higher SFC), and the same trend was also found in the PCA results (Figure 7 A and B).

### 3.4 Individual identification

After analyzing the informational difference and the methodological difference at the predicted-connectivity level and SFC level, we utilized the SFC calculated by each approach in Table 1 for individual identification to explore the impact of the informational difference and the methodological difference in the specific scenario of individual identification. We used different classifiers and the number of subjects to identify individuals to ensure the reliability of the conclusions. The accuracy of individual identification is shown in Figure 8. As shown in the figure, for the regression approach and the MLP, the SFC calculated by the direct model and the full model are almost identical in terms of individual identification, which indicates that their accuracy of individual identification is mainly affected by the methodological difference. This is consistent with the results of the difference analysis at SFC level. However, for the pGCN, the SFC calculated by the direct model and the full model differs obviously in individual identification, even greater than the methodological difference in some cases (i.e., when the violin of the correlational approach lies between the violins of the direct model and the full model). Furthermore, the informational difference has different effects under different prediction directions. The informational difference in the SFC calculated by the pGCN in the SC-to-FC direction reduces the identification accuracy, while in the FC-to-SC direction, it may increase the identification accuracy. However, the opposite occurred in the sGCN. Compared to the SFC calculated by the direct sGCN, the SFC calculated by the full-SC sGCN shows higher accuracy in individual identification, but the SFC calculated by the full-FC sGCN shows lower accuracy. However, similar to the pGCN, the informational difference also has a greater impact on the identification accuracy of the SFC calculated by the sGCN in some cases.

**Figure 8:**
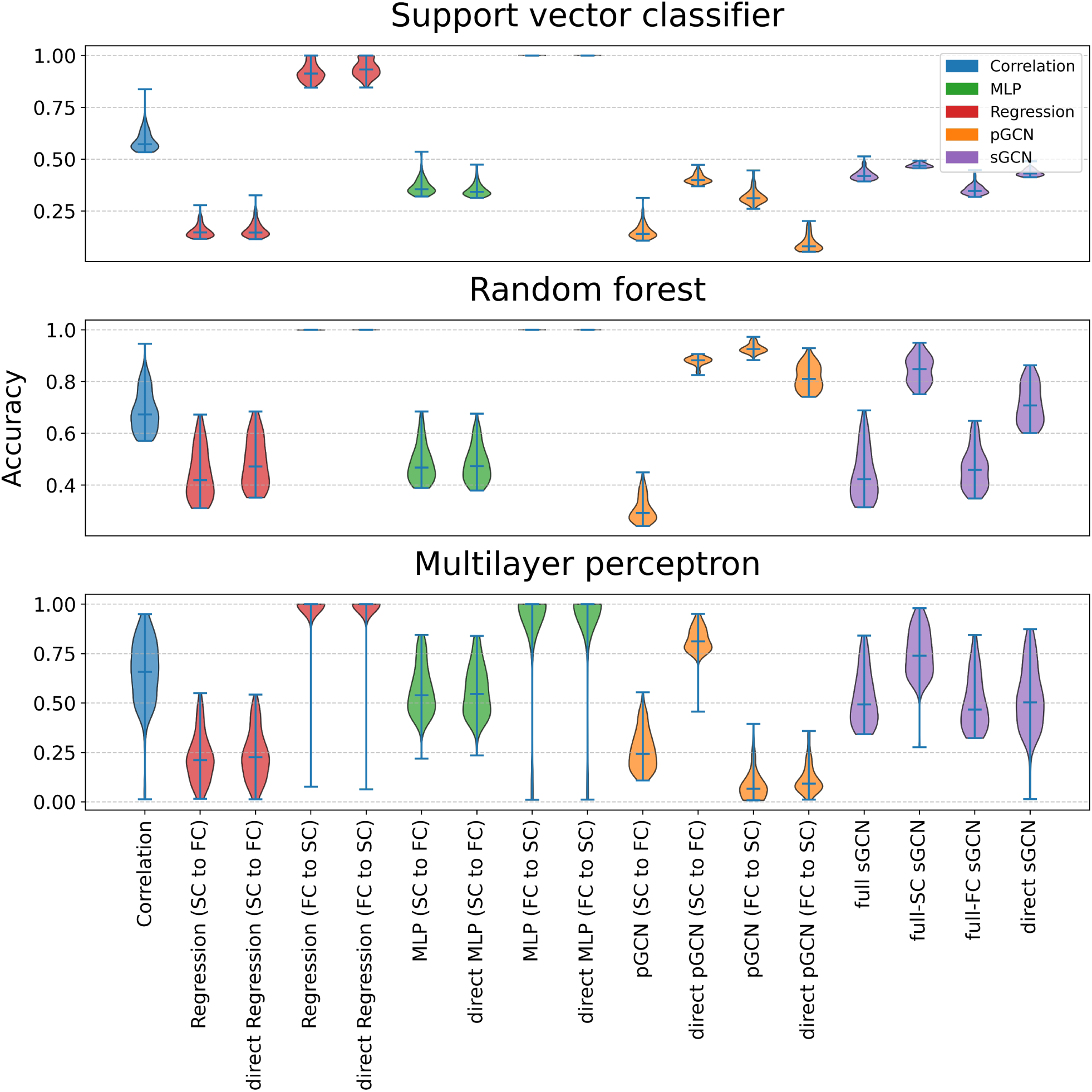
Accuracy of individual identification using different classifiers and SFCs. Each violin represents the accuracy distribution of individual identification using the SFC calculated by each model across different numbers of subjects (from 10 to 100). Violins with the same color represent that the SFCs they used come from the same model, but with different combinations of information.

## 4 Discussion

In this study, we used five approaches to evaluate structure-function coupling (SFC) and compared the informational difference and the methodological difference. Three were predictive approaches (Figure 4 A): regression approach (Vázquez-Rodríguez et al., 2019), multilayer perceptron (MLP) (Sarwar et al., 2021), and predictive graph convolutional network (pGCN) (Chen et al., 2024). The other two were non-predictive approaches (Figure 4 B): correlational approach (Baum et al., 2020; Gu et al., 2021; Honey et al., 2009; Liégeois et al., 2020) and self-supervised graph convolutional network (sGCN). We first demonstrated that all five approaches captured effective coupling patterns, then analyzed the differences at the predicted-connectivity level and SFC level, and finally examined the effect of these two differences on individual identification. Our study shows that the informational difference has a small effect on the regression approach and the MLP, but a larger effect on the pGCN and the sGCN. Furthermore, we observed that, at the predicted-connectivity level, the absolute value of the informational difference is positively correlated with myelination index when predicting functional connectivity (FC), while at SFC level, the informational difference is lowest in the right-hemispheric dorsal attention network and highest in the orbito-affective network. Our study contributes to understanding the sources and manifestations of the differences between modeling approaches and the correlational approach when evaluating SFC, and the role of indirect connections in SFC calculation.

Quantifying the correspondence between structural and functional connectivity is an important issue in neuroscience (Damoiseaux & Greicius, 2009; Huang & Ding, 2016; Wang et al., 2015), but different approaches often yield different degrees of SFC. For example, traditional correlational approaches and computational models often yield a lower SFC, while deep learning models often yield a higher SFC (Chen et al., 2024; Esfahlani et al., 2022; Sarwar et al., 2021; Vázquez-Rodríguez et al., 2019). More importantly, these approaches also differ in their evaluations of the relative strength of the regional SFC. This is not only related to individual differences and different preprocessing procedures, but also reflects the systematic differences in SFC calculation among these approaches. In this study, we hypothesize that the differences in the information combination used by different approaches belong to this systematic difference, which we define as informational difference, while the differences that still exist when using the same information combination are defined as methodological difference. Therefore, based on the three typical predictive models we reproduced, as well as the self-supervised model we designed and the traditional correlational approach, we systematically analyzed these two differences.

Analysis at the three levels of predicted-connectivity, structure-function coupling, and individual identification all indicate that the informational difference has little effect on the regression approach and the MLP, but a greater effect on the pGCN and the sGCN. In other words, the introduction of indirect connectivity has little effect on the regression approach and the MLP when capturing the coupling relationship between SC and FC, but plays a more important role in GCN models. Previous studies have shown that indirect connections help improve the predictive ability for FC (Esfahlani et al., 2022; Røge et al., 2017), which is confirmed in our reproduced predictive GCN model (Figure 6 C). However, there may still exist predictive models that are insensitive to indirect connections, such as the multiple linear regression and the MLP that we reproduced. In this study, we even found that, in some regions, the introduction of indirect connections may increase the model’s prediction error (e.g., Figure 6 B, the MLP’s prediction of contralateral homologous). Furthermore, we found that the insensitivity of the regression approach and the MLP to indirect connections is reflected in both the SC-to-FC and FC-to-SC prediction directions, further increasing the reliability of our conclusions.

Regarding the specific distribution of the informational difference, we found that at the predicted-connectivity level, the absolute informational differences of the three predictive models are positively correlated with myelination index (Figure 6 G-I). This means that the prediction of direct connections in cortical regions with higher myelination indices is more significantly affected by indirect connections. Previous studies have shown that T1w/T2w myelination index gradually decreases along the sensorimotor-association axis from the primary sensorimotor cortex to the association cortex (Baum et al., 2022; M. F. Glasser & Essen, 2011). Therefore, our results suggest that the primary sensorimotor cortex is more affected by indirect connections when predicting FC, while the association cortex is less affected. We speculate that this may be because the primary sensorimotor cortex has stronger structure-function coupling (Collins et al., 2024; Vázquez-Rodríguez et al., 2019), thus capturing stable structure-function correspondences solely through direct connections. Therefore, the introduction of indirect connections can lead to larger fluctuations in the correspondences, resulting in fluctuations in the prediction error, making it more sensitive to indirect connections. The opposite is true for the association cortex. It should be noted that the correlation between the informational difference and myelination index appears only stably in the SC-to-FC prediction direction. This may be because myelination index reflects structural properties instead of functional properties, this point is also observed in the Pearson correlation coefficient between SFC and myelination index in Table 2. At SFC level, we found that the informational difference was lowest in the right-hemispheric dorsal attention network and highest in the orbito-affective network. This conclusion was confirmed in both the SC-to-FC and FC-to-SC prediction directions (Figure 7 A). It indicates that the SFC of the right-hemispheric dorsal attention network is less affected by indirect connections, while the SFC of the orbito-affective network is more significantly affected by indirect connections. We speculate that this may be related to the hemispheric asymmetries of the dorsal attention network (Alam et al., 2022; Duecker & Sack, 2015) and the difference in connectivity between the orbito-affective and dorsal attention networks (Spreng et al., 2013; Zald et al., 2012).

This study has several limitations. First, our analysis was conducted on only one dataset, making the conclusions susceptible to influences from specific data sources and preprocessing procedures, thus reducing their generalizability. To address this, we considered two prediction directions (SC-to-FC and FC-to-SC) as well as multiple predictive models, whose consistent performance on the main conclusions enhances their robustness. Second, in this study, we used a 50% threshold to sparsify the connectivity matrices, this is an empirical threshold that may filter out weak but important connectivity signals. Finally, the predictive performance of the multiple linear regression model and its susceptibility to indirect connections may be related to the choice of independent variables, we only considered the three structural metrics used by Vázquez-Rodríguez et al. (2019), which may have limitations.

Despite these limitations, our study shows that indirect connections have varying effects on different models and modalities. Indirect connections have a smaller effect on the regression approach and the multilayer perceptrons, but a larger effect on GCN models. Specifically, at the predicted-connectivity level, cortical regions with higher T1w/T2w myelination indices are more significantly affected by indirect connections in predicting direct connections. At structure-function coupling level, indirect connections have the least effect on the right-hemispheric dorsal attention network, but the greatest effect on the orbito-affective network. Furthermore, our study shows that indirect connections do not always have a positive effect, they can also increase prediction error in connectivity and decrease the individual identification ability of structure-function coupling. In summary, our study helps to understand the sources and components of the differences between modeling approaches and the correlational approach in the calculation of structure-function coupling, and suggests that the selection of the approach for calculating structure-function coupling in future studies that use structure-function coupling as biomarkers should be more careful.

## Data and Code Availability

The original structural connectivity and the parcellation templates come from https://doi.org/10.5281/ zenodo.10150880, and the functional connectivity data come from https://db.humanconnectome.org. The preprocessed structural and functional connectivity, as well as other data such as structure-function coupling in this study, are available at https://doi.org/10.5281/zenodo.19622237. The preprocessing and modeling code can be found in https://github.com/ZhYhn/SFC difference.

## Author Contributions

Y.Z: Conceptualization, Data curation, Formal analysis, Investigation, Methodology, Software, Validation, Visualization, Writing—original draft, Writing—review and editing.

## Acknowledgements

Data were provided by the Human Connectome Project, WU-Minn Consortium (Principal Investigators: David Van Essen and Kamil Ugurbil; 1U54MH091657) funded by the 16 NIH Institutes and Centers that support the NIH Blueprint for Neuroscience Research; and by the McDonnell Center for Systems Neuroscience at Washington University.

## Supplementary Material

Supplementary material for this article is available with the online version here: https://doi.org/10.5281/zenodo.19622237

