## Supplementary figures for "Informational and methodological differences in regional structure-function coupling in modeling approaches"

**Figure S1.** Structure-function coupling calculated by five approaches. **A**-**E** Mean structure-function coupling (SFC) of the test set calculated by five approaches. **F** The Pearson correlation matrix of the five SFCs.

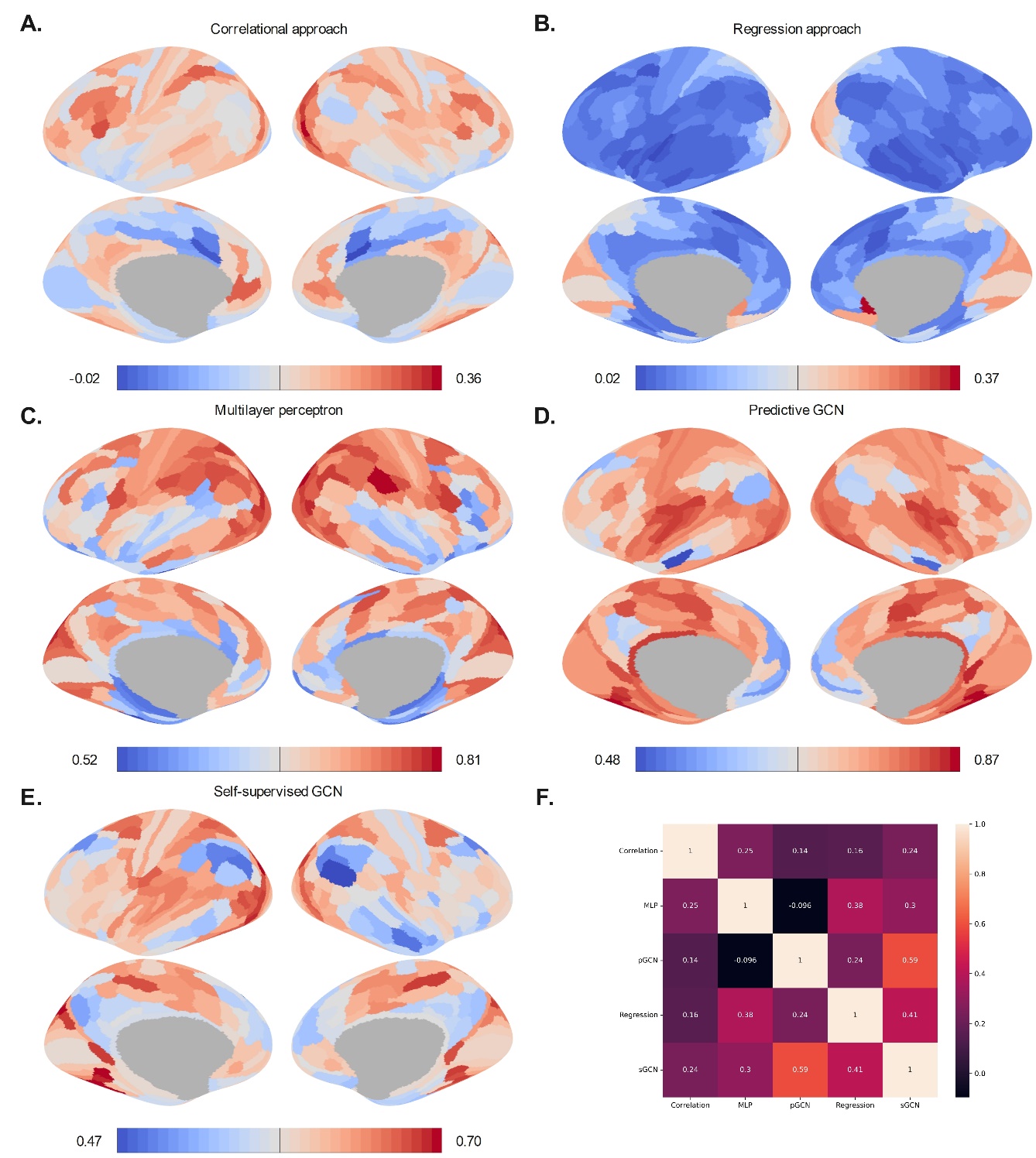

**Figure S2.** The informational difference and the methodological difference at the predicted-connectivity level. **A** The proportions of positive and negative values in the informational difference matrix and the methodological difference matrix. **B** The means of positive and negative values in the informational difference matrix and the methodological difference matrix.

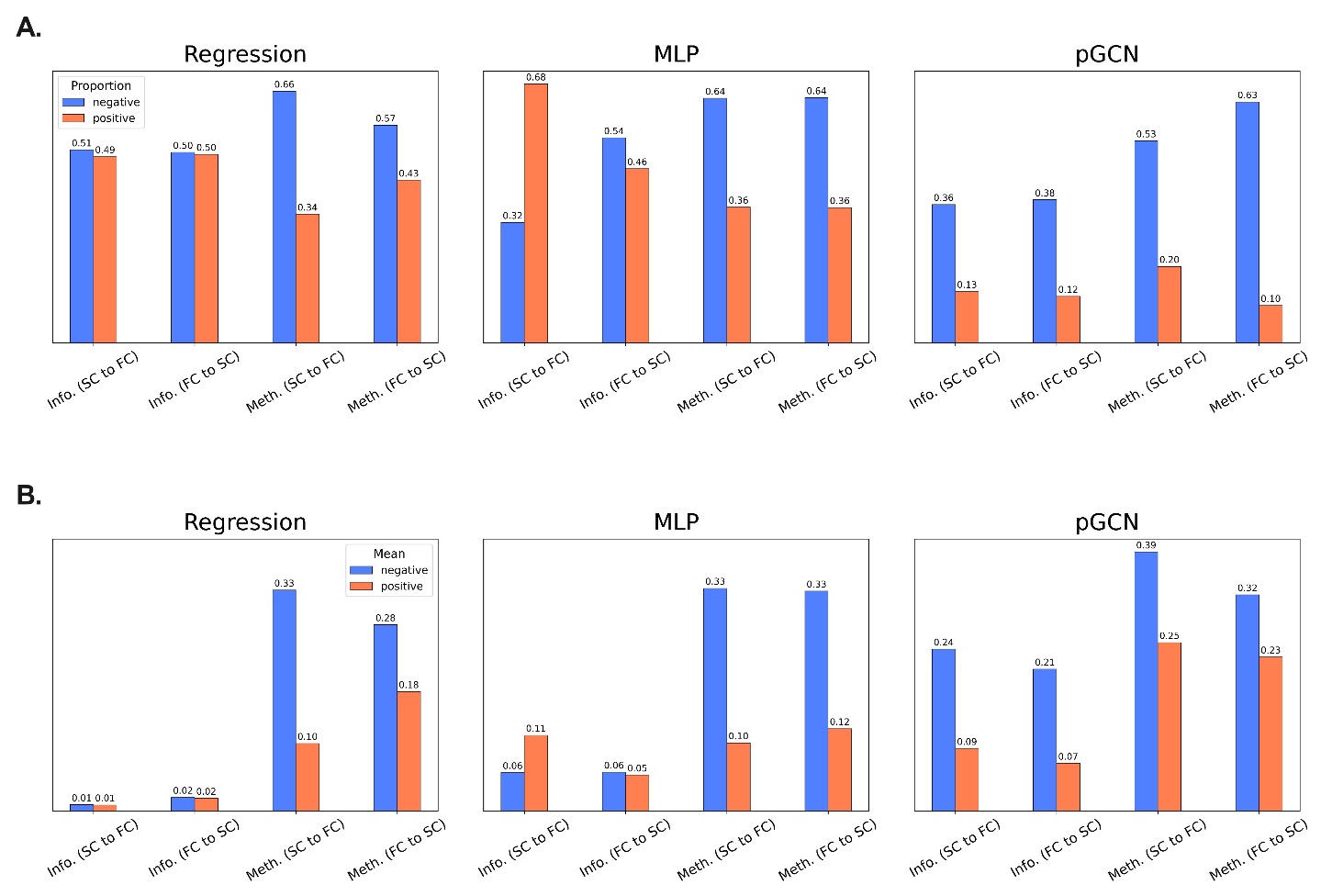
**Figure S3.** The informational difference and the methodological difference at SFC level. **A-Q** Mean differences of the
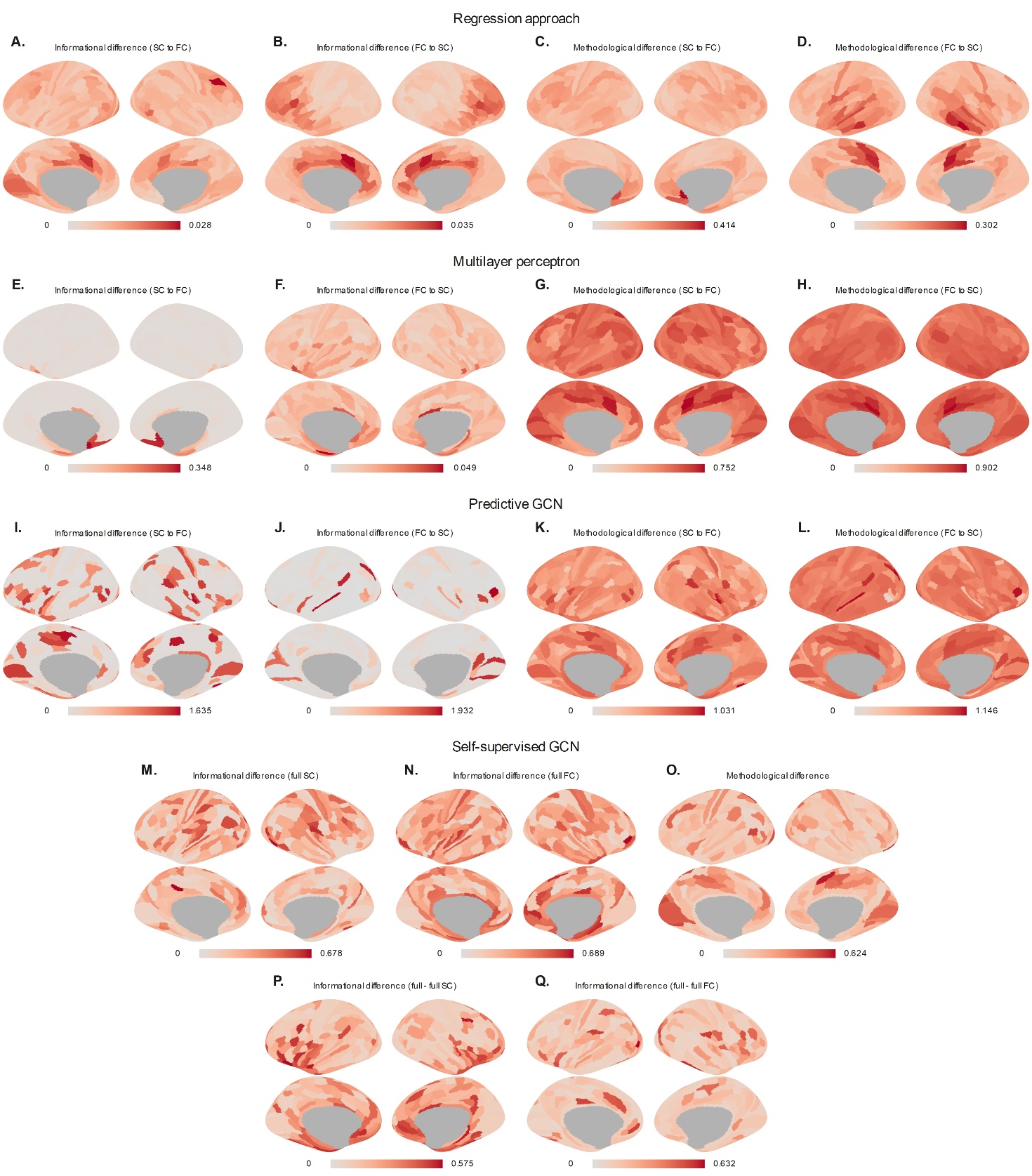
test set calculated by the five approaches.
